# Meiotic prophase I disruption as a strategy for non-hormonal male contraception using small-molecule inhibitor JQ1

**DOI:** 10.1101/2025.05.13.653822

**Authors:** Leah E. Simon, Stephanie Tanis, Adriana K. Alexander, Tegan Horan, Mercedes Carro, Samantha-Jane Bonnett, Roni Ben-Shlomo, Audrey Xie, Charles G. Danko, Jelena Lujic, Paula E. Cohen

**Author notes:** Contributed equally (author order selected randomly) (Leah E. Simon, Stephanie Tanis). **Corresponding author:** Paula E. Cohen.

## Abstract

Long-acting non-hormonal male contraceptives are urgently needed but developing strategies that are both effective and reversible presents significant challenges. Here, we investigated the potential of meiotic prophase I blockade as a promising and potentially reversible approach to male contraception. To do this, we utilized (+)-JQ1, a small-molecule inhibitor of the testis-specific protein, BRDT. Daily injections of (+)-JQ1 for three weeks resulted in disrupted spermatogenesis resulting in loss of spermatozoa and an inability to sire pups. While spermatogenic cells repopulated the testis within six weeks post drug cessation, full fertility restoration required a longer recovery period. We attribute this delay in full recovery to persistent issues with the pachytene transcriptional program, which is crucial for meiotic progression and spermatid development. These findings underscore the potential of pharmacological approaches to disrupt meiotic prophase I as a targeted, reversible male contraceptive strategy, providing new insights into developing effective non-hormonal contraceptive approaches.

## INTRODUCTION

Unintended pregnancies account for nearly 44% of all pregnancies worldwide, highlighting the urgent need for expanded contraceptive options, particularly male-targeted methods (Finer and Zolna 2016; Bearak et al. 2020). Despite this demand, most available contraceptives either rely on female-centered approaches or require interventions such as vasectomy with potentially permanent and irreversible effects, leaving a significant gap in reversible male contraception (Castaneda and Matzuk 2015). Developing non-hormonal strategies that temporarily and specifically suppress sperm production would help address this imbalance. One promising approach is targeting key regulators of spermatogenesis, the multi-step process by which diploid germ cells divide and differentiate into mature spermatozoa. This process, which continues throughout adulthood, involves, through broadly defined genetic programs, the mitotic proliferation of spermatogonia, meiotic division of diploid spermatocytes into haploid spermatids, and finally post-meiotic differentiation of spermatids to become mature spermatozoa (Griswold 2016; Fayomi and Orwig 2018). By selectively disrupting critical regulators at specific stages, it may be possible to reversibly impair sperm production without permanently affecting fertility. Advances in genomics and single-cell reference datasets now allow us to identify the specific genetic program that drives each stage of spermatogenesis, and thereby to pinpoint precise developmental windows that may be amenable for intervention (Green et al. 2018; Hermann et al. 2018; Ernst et al. 2019; Grive et al. 2019). This unprecedented level of target specificity opens the door to develop non-hormonal male contraceptives that act at well-defined molecular targets without compromising long-term fertility.

Selecting the optimal developmental window for intervention remains a key challenge in contraceptive development. Early-stage disruptions targeting spermatogonia might risk impairing the stem cell niche, raising concerns about long-term fertility restoration. Conversely, post-meiotic interventions may allow some sperm to escape, leading to potential fertilization. An ideal contraceptive strategy would therefore act at a stage where sperm production is effectively halted while ensuring full recovery upon cessation of treatment. Meiosis presents as an attractive target in this regard, as it represents a bottleneck in sperm production where precise regulatory mechanisms control the transition from diploid precursors to haploid gametes.

Among the meiotic stages, prophase I stands out as a particularly promising point of intervention. This transcriptionally active phase is critical for homologous chromosome pairing, synapsis, and recombination, making it essential for the progression of spermatogenesis (Handel and Schimenti 2010; Gray and Cohen 2016). Disrupting key regulators at this stage could have pronounced effects on spermatogenesis while avoiding the risks associated with stem cell depletion or incomplete post-meiotic inhibition. Moreover, because the transcriptional program established in pachytene (mid-prophase I) spermatocytes largely determines later sperm development (Alexander et al. 2023; Kaye et al. 2024), interventions at this stage could exert sustained contraceptive effects without requiring prolonged drug exposure. Conversely, a major concern is whether transient pharmacological interference during meiosis could lead to persistent defects, such as aneuploidy, chromosomal mis-segregation, or long-term transcriptional alterations that impact gamete function and ultimately fertility. This challenge is further compounded by the blood-testis barrier (BTB), which limits pharmaceutical access to the seminiferous tubules and target cells like meiotic spermatocytes (Yan Cheng and Mruk 2012). Matzuk et al. previously examined the contraceptive effects of (+)-JQ1 (JQ1), a small molecule inhibitor that competitively binds to the bromodomains of BRDT, a testis-specific transcription factor that is essential for meiotic prophase I progression (Shang et al. 2004; Shang et al. 2007; Dhar et al. 2012; Matzuk et al. 2012). While their landmark study detailed the impact of JQ1 on several reproductive parameters, the cytological, molecular and transcriptomic effects of meiotic disruption and recovery after drug withdrawal remain unexplored. Given the critical transcriptional burst observed during pachynema of prophase I, which regulates genes essential for spermiogenesis, a deeper investigation is needed to assess the potential of meiotic prophase I as a fully reversible, non-hormonal contraceptive target while ensuring long-term reproductive safety.

Here, we investigated the effects of pharmacological disruption of meiotic prophase I using JQ1. Given the key role of BRDT in coordinating the transcriptional burst during pachynema of prophase I (Alexander et al. 2023), JQ1 targeting provides an opportunity to assess whether transient inhibition at this stage induces lasting disruption in spermatogenesis. Male mice were treated for three weeks with JQ1, followed by assessment of fertility, gene expression, and prophase I progression using single cell RNA sequencing (scRNA-seq), histology, and chromosome spreading. To evaluate reversibility, we examined recovery of fertility parameters at 6- and 30-weeks post-treatment, and we also assessed reproductive outcomes in the F1 progeny. Our findings demonstrate that JQ1 treatment results in altered transcription dynamics in spermatocytes, leading to a loss of post-meiotic cells. While these spermatogenic cells repopulated within six weeks post drug cessation, full fertility restoration and recombination events during prophase I was restored following 30 weeks post drug cessation. Furthermore, transient meiotic inhibition had no impact on the fertility of subsequent generations. These findings support meiotic prophase I as a promising target for reversible male contraception, alleviating concerns about permanent disruptions to fertility. Crucially, this study is the first to comprehensively chart the recovery of spermatogenesis across multiple post-treatment time points, demonstrating not only the robustness but also the timeline of fertility restoration. By pinpointing this specific stage in spermatogenesis, and by demonstrating robust reversibility of the contraceptive downregulation of meiosis prophase I, this study provides a framework for future investigations into pharmacological inhibition of meiosis as a viable male contraceptive strategy.

## RESULTS

### JQ1 treatment leads to the depletion of post-meiotic spermatids, with full recovery occurring after drug withdrawal

Despite advances in understanding the molecular regulation of spermatogenesis through gene targeting approaches in the laboratory mouse, *Mus musculus*, the consequences of transient pharmacological disruption of meiosis and the timeline of recovery remain poorly understood. Thus, to investigate effects of BRDT inhibition on meiotic prophase I dynamics and molecular regulation, as well as the impact on these processes following drug withdrawal, we treated sexually mature DBA/2J male mice with daily intraperitoneal (i.p.) injections of JQ1 (50 mg/kg; treatment group, T) or vehicle control (vehicle control-treatment, VC-T) for three weeks. This treatment duration has been shown to induce a contraceptive effect by disrupting spermatogenesis during prophase I of meiosis (Matzuk et al. 2012). After treatment, half of each cohort was sacrificed for analysis. The remaining animals (recovery, R, or vehicle control-recovery, VC-R) were examined six weeks post-treatment to assess the effects of meiotic resumption on fertility and spermatogenesis post drug cessation. Age-matched DBA/2J males were included as untreated controls for both treatment (UC-T) and recovery (UC-R) phases (Figure 1A).

**Figure 1.**
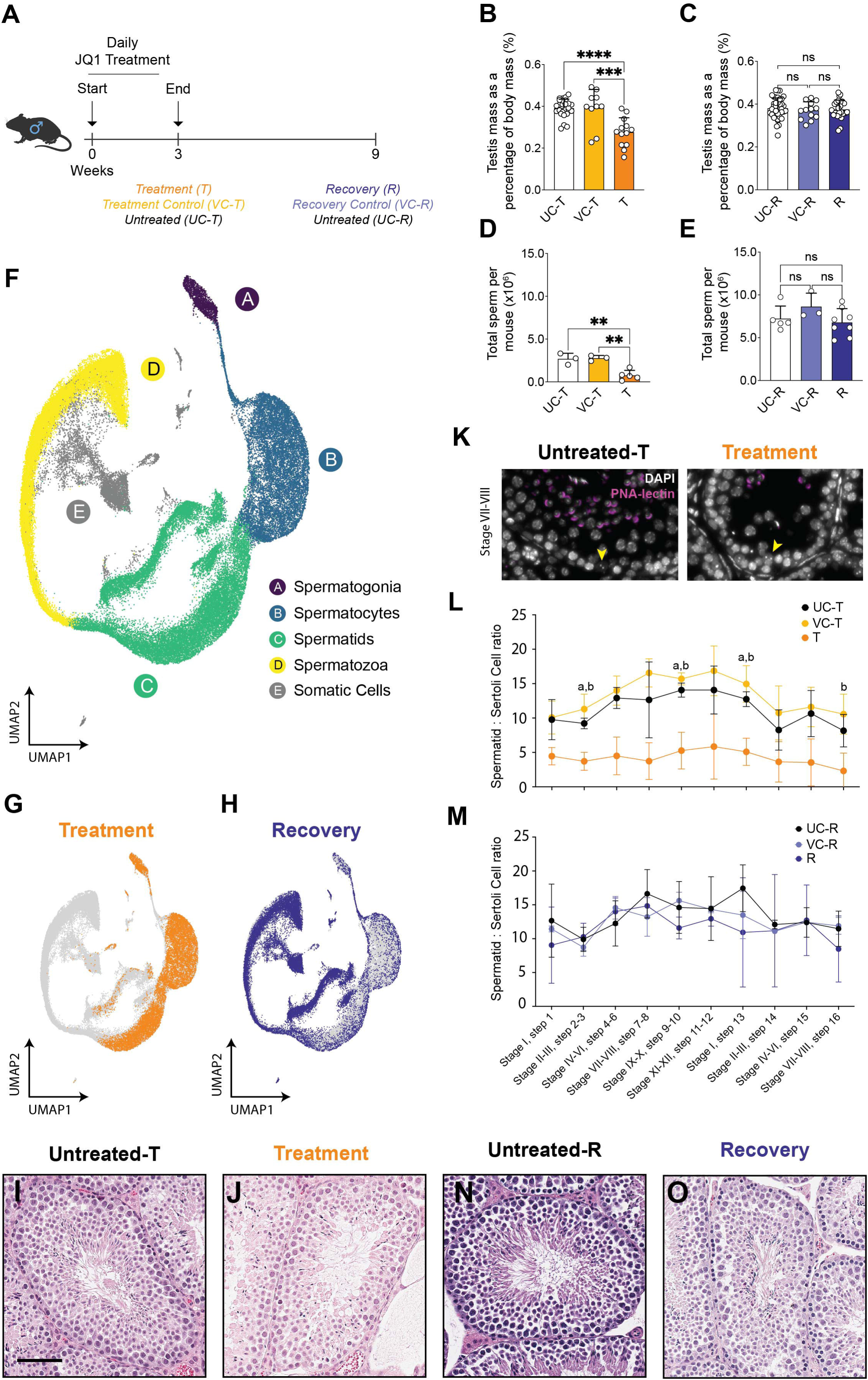
Three-week JQ1 treatment results in post-meiotic cell loss. (A) Experimental design outlining three-week daily JQ1- (T) or vehicle injection (VC-T) with age-matched untreated controls (UC-T), and recovery six weeks post JQ1- (R) or vehicle (VC-R) cessation with age-matched untreated controls (UC-R). (B, C) Testis mass per mouse as percentage of body mass (% ± SD) for (B) T (*n* = 8), VC-T (*n* = 5), UC-T (*n* = 13), and (C) R (*n* = 13), VC-R (*n* = 7), and UC-R (*n* = 20) mice. (D, E) Total epididymal sperm counts (sperm per million in ± SD) for (D) T (*n* = 5), VC-T (*n* = 5), UC-T (*n* = 3) mice, and (E) in R (*n* = 8), VC-R (*n* = 3), UC-R (*n* = 5) mice. (F) UMAP visualization representing all cells captured for all scRNA-seq Treatment and Recovery timepoints (*n* = 12) and untreated control (*n* = 3) libraries. (G) UMAP visualization of cells captured following Treatment. (H) UMAP visualization of cells captured following Recovery. (I, J) Testicular histology for Untreated-T (I) and Treatment (J) as shown by H&E staining of paraffin embedded testis sections. (K) Staining of testis sections at stage VII-VIII using PNA-lectin (magenta) and DAPI (grayscale) from Untreated-T and Treatment animals. Yellow arrows indicate Sertoli cells. (L, M) Ratio of spermatids relative to Sertoli cells per tubule of specific stages in (L) UC-T, VC-T, and T (*n* = 3 mice) and (M) in VC-R and R (*n* = 2 for both), and UC-R (*n* = 3) mice (mean ± SD). ^a^ p < 0.05 T compared to UC-T, ^b^ p <0.05 T compared to VC-T. (N, O) Testicular histology for Untreated-R (N) and Recovery (O) as shown by H&E staining. Scale bar = 100 µm. P values obtained from a one-way ANOVA test (ns = not significant, *p < 0.05, **p<0.01, ***p<0.001, ****p<0.0001).

To confirm the effectiveness of our treatment and recovery timepoints, we first evaluated general reproductive parameters (Matzuk et al. 2012). JQ1-treated animals showed significant reductions in testis mass (Figure 1B) and sperm counts (Figure 1D) compared to both VC-T and UC-T. However, by six weeks post-treatment, testis mass (Figure 1C) and sperm counts (Figure 1E) of R animals had returned to baseline levels, comparable to VC-R and UC-R animals.

Next, we examined the gene expression dynamics underlying JQ1-induced germ cell loss and recovery. Single-cell RNA sequencing (scRNA-seq) of testicular suspensions from T and R timepoints, along with vehicle-treated and untreated controls (VC-T, VC-R, UC-T, UC-R), provided a high-resolution view of germ cell dynamics. Following pre-processing of scRNA-seq data (see Methods; Supplementary Figures S1-3), we analyzed 69,262 cells from the testes of all experimental groups, capturing major germ and somatic cell types. Clustering analysis identified four major germ cell types, visualized using Uniform Manifold Approximation and Projection (UMAP) (Figure 1F), consistent with prior studies on mouse spermatogenesis (Green et al. 2018; Hermann et al. 2018; Lukassen et al. 2018; Grive et al. 2019).

Single cell RNA-seq analysis revealed that cells progress to the haploid spermatid stage of spermatogenesis but fail to develop into spermatozoa (Figure 1G), supporting the contraceptive effect of JQ1. This was further validated by hematoxylin and eosin (H&E) staining of paraffin embedded testicular sections which revealed fewer spermatozoa in T males relative to UC-T males (Figure 1I, J). Using PNA-lectin histochemistry staining, which binds post-meiotic cells (Cheng et al. 1996), we further observed that spermatids, though present, were reduced across all steps (1–16) in T males compared to controls (Figure 1K, L). Finally, TUNEL staining indicated a trend in increased apoptosis in T males relative to UC-T and VC-T males; however, this trend was not significant (Supplementary Figure S4A). Together, these results demonstrate that JQ1 treatment disrupts spermatid development and supports its potential as a contraceptive by depleting both spermatid and spermatozoa populations.

Six weeks after JQ1 withdrawal, we observed full recovery of spermatogenesis, with all major germ cell types reappearing in libraries from R germ cells, as indicated by the UMAP (Figure 1H) and H&E staining of testis sections (Figure 1N-O). This repopulation of germ cells within the testis with PNA-lectin-positive haploid cells was further confirmed by lectin histochemistry staining of testicular sections and quantification of spermatid steps, where all spermatid steps (1-16) were comparable across all conditions (Figure 1M). Furthermore, TUNEL staining indicated no difference in apoptotic cells across all condition groups (Supplementary Figure S4B). Collectively, these results demonstrate that JQ1 treatment induces significant loss of spermatids and spermatozoa. Importantly, these cell populations were fully restored within one spermatogenic cycle following cessation of JQ1, establishing reversibility of the contraceptive effect within six weeks.

### Haploid cell loss is driven by JQ1-induced disruption of the meiotic transcriptional burst

To determine the timing of haploid cell loss, we performed pseudotime analysis on scRNA-seq libraries using Slingshot (Street et al. 2018) (Supplementary Figure S5). Cells were grouped into 51 bins along the spermatogenesis timescale, revealing loss specifically in T libraries during the spermatid-to-spermatozoa transition (Figure 2A; dashed line). To assess whether this loss was linked to transcriptional changes, we analyzed testicular cells from UC-T, VC-T, and T, plotting transcript diversity along pseudotime. Control samples exhibited a characteristic transcriptional burst in spermatocytes followed by a smaller burst in spermatids, which declined during late spermiogenesis (Alexander et al. 2023) (Figure 2B). In contrast, JQ1-treated spermatocytes showed reduced transcriptional diversity (Figure 2B; dashed rectangle), indicating disruption of the meiotic transcriptional burst. It remains to be explored whether this disruption in the transcriptional burst also affects the integrity of meiotic prophase I.

**Figure 2.**
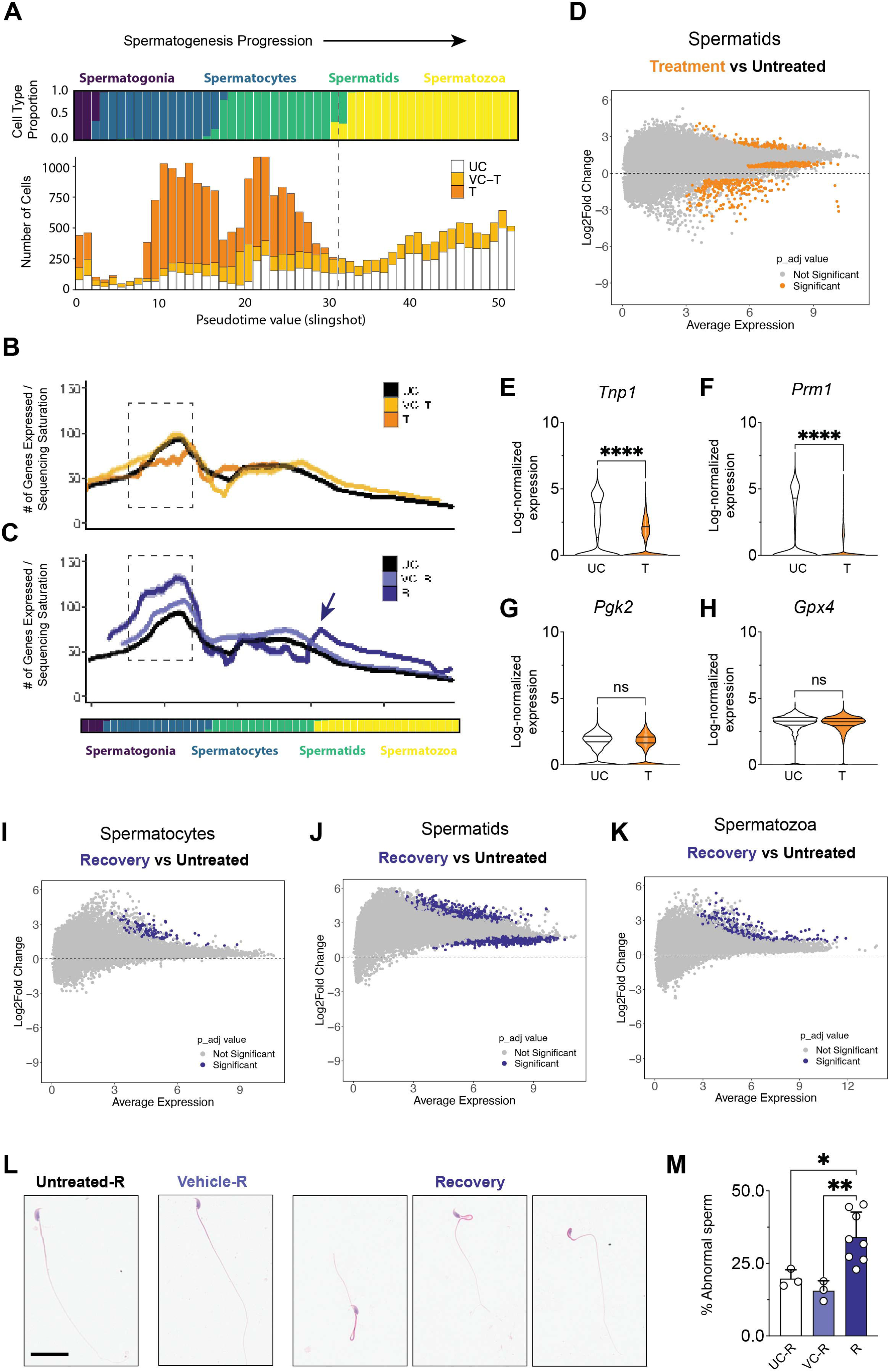
Haploid cell loss is driven by transcriptional changes that do not recover. (A) Pseudotime analysis on scRNA-seq libraries using Slingshot showing spermatogenesis progression (top panel) and number of cells per condition group (bottom panel). Cells are grouped into 51 bins along the spermatogenesis timescale. Dashed line indicates the spermatid-to-spermatozoa transition. (B, C) Transcript diversity plotted along pseudotime for (B) UC-T, VC-T, and T and (C) UC-R, VC-R, and R (dashed box shows spermatocytes). Purple arrow indicates the spermatid-to-spermatozoa transition. (D) DE analysis on pseudobulked spermatids from scRNA-seq identifying 518 differentially expressed genes between Treatment and Untreated control groups. (E, F) Examples of differentially expressed genes between T and UC groups. (G, H) Examples of genes that are not differentially expressed between T and UC. Violin plots show gene expression in individual cells, grouped by condition. Statistical comparisons are based on average expression per biological replicate (*n* = 3 per group). P values obtained from a Student’s t-test (ns = not significant, ****p<0.0001). (I-K) DE analysis on pseudobulked spermatocytes showing 186 genes (I), spermatids showing 533 genes (J), and spermatozoa showing 308 genes (K) from scRNA-seq between R and UC groups. (L) Representative epididymal sperm, stained with H&E, from Untreated-R, Vehicle-R and Recovery males, highlighting major abnormalities. (M) Quantification of sperm abnormalities (mean ± SD) from UC-R (*n* = 3), VC-R (*n* = 5) and R (*n* = 5) males. P values obtained from a one-way ANOVA test (ns = not significant, *p < 0.05, **p<0.01).

To confirm that JQ1 acts via competitive inhibition of BRDT rather than through transcriptional repression, we assessed RNA expression and protein localization. *Brdt* mRNA levels remained unchanged in T animals (Supplementary Figure S6A), while BRDT protein remained present in the testis (Supplementary Figure S6B), supporting the role of JQ1 as a competitive inhibitor rather than a transcriptional repressor.

During pachynema of prophase I, spermatocytes undergo a transcriptional burst, activating numerous genes essential for both meiosis and spermiogenesis. While meiosis-related genes peak before pachynema, the expression of late spermatogenesis genes surges at the zygotene-to-pachytene transition, ensuring proper progression of post-meiotic processes (Geisinger et al. 2021). Disruption of the prophase I transcriptional burst may compromise key regulatory programs required for spermatid survival, ultimately leading to their loss following JQ1 treatment (Alexander et al. 2023; Carro et al. 2025). To investigate whether downstream transcriptional changes contribute to this depletion, we performed differential expression (DE) analysis on pseudobulked reads from our scRNA-seq dataset of spermatids from treated and untreated mice. This analysis identified 518 differentially expressed genes between T and UC groups (Figure 2D). Many of the most downregulated genes in the T group have established roles in sperm formation and maturation, including *Tex44* (Dupuis et al. 2024), *Spem2* (Li et al. 2024), and *Tssk2* (Xu et al. 2008), as well as chromatin remodeling during the later stages of spermiogenesis, including *Smarca5* (Kataruka et al. 2024), *Tnp1* (Zhao et al. 2004), *Tnp2* (Zhao et al. 2001) (Figure 2E), *Prm1* (Cho et al. 2001) (Figure 2F), and *Prm2* (Cho et al. 2003). Decreased accessibility of several of these genes has been reported in JQ1-treated spermatids (Wang et al. 2022), which may contribute to the transcriptional repression observed here. Moreover, recent ChIP-seq studies profiling BRDT occupancy have shown that in addition to wide-spread BRDT binding in the mouse genome, binding sites of BRDT are mainly located in gene units such as promoters, introns and exons, and that BRDT cooperates with specific TFs to regulate genes in meiotic and post-meiotic cells (Her et al. 2021). Our results showed that genes essential for spermiogenesis that are not bound by BRDT showed no expression changes (Figure 2G, H), demonstrating that JQ1 treatment does not induce a broader effect on transcriptional activity genome-wide, but is instead largely restricted to genes whose promoters contain BRDT binding motifs (Imai et al. 2009; Danshina et al. 2010; Her et al. 2021). Collectively, these findings demonstrate that disruption of the pachytene transcriptional burst impairs essential regulatory networks required for spermatid survival, contributing to their depletion.

### Repopulating testicular cells exhibit persistent transcriptional changes

To assess the restoration of the spermatogenic program following JQ1 withdrawal, we examined the transcriptional profiles of recovery libraries from untreated, vehicle-treated and treated recovery groups (UC, VC-R, and R). This analysis provides critical insights into the extent to which meiosis-targeted contraception is reversible and whether persistent transcriptional changes could impact future fertility or gamete quality/viability. Along the pseudotime axis, we observed that while spermatocytes in R libraries regained expected transcriptional patterns, including a coordinated burst of transcriptional diversity (Figure 2C; dashed rectangle), the magnitude of this burst exceeded that observed in VC-R or UC (Figure 2C). This restored burst of transcriptional activity was further supported by DE analysis of pseudobulked expression data from spermatocytes post-JQ1 cessation, which identified upregulation of 156 of a total 186 genes in R libraries compared to that of the UC group (Figure 2I), potentially as a compensatory response to prior inhibition.

While VC-R and UC controls displayed the characteristic smaller burst in expression at pachynema followed by a tapering off of transcriptional activity in spermatids, DE analysis revealed that spermatids from R libraries exhibited widespread dysregulation, with 224 downregulated genes and 309 upregulated genes (Figure 2J). This suggests that transcriptional homeostasis is not fully restored in repopulating spermatids following drug withdrawal. In spermatozoa, the transcript diversity plot revealed a pronounced increase in transcriptional activity at the transition to spermatozoa, which persisted throughout the remainder of the pseudotime axis (Figure 2C; arrow). DE analysis showed a total of 308 genes, all of which were upregulated, a predominance of upregulated genes, indicating a failure to fully re-establish the usual critical transcriptional silencing mechanisms in repopulating spermatozoa following drug withdrawal (Figure 2K).

Given that transcriptional misregulation, particularly in the later stages of spermatogenesis, has been linked to morphological defects in spermatozoa (Wang et al. 2019; Moritz and Hammoud 2022), we examined epididymal spermatozoa in post-treatment recovery (R) mice. Compared to UC-R and VC-R controls, R mice exhibited a significant increase in epididymal spermatozoa exhibiting morphological defects (19.72% ± 3.06 for UC-R; 15.58% ± 3.44 for VC-R; 34.05% ± 8.65 for R; p<0.05) (Figure 2L, M). These findings demonstrate that while spermatogenic cells repopulate the testes after JQ1 withdrawal, unresolved transcriptional deficiencies may impact the repopulating gametes. This highlights the potential long-term effects of meiosis-targeted contraception and underscores the need for further investigation into how these transcriptional irregularities may impact fertility.

### Fertility and sperm quality take longer to recover following resumption of spermatogenesis

Since the ultimate test of any potential contraceptive drug is the ability to prevent pregnancy without impacting virility or male sexual behavior, we next asked whether JQ1-treated males would mate with cycling female mice, and whether such mating events could result in the production of viable offspring. Following 3 weeks of daily JQ1- or vehicle injections, males were placed in a breeding assay with mature DBA/2J adult females on a biweekly rotating schedule, wherein a female was placed with a male for two weeks and replaced with another female. Age matched untreated males were also placed in the same breeding assay as controls. Although spermatogenesis resumption occurred within six weeks following JQ1 withdrawal, we observed a significant delay in the time to first litter for T males compared to VC-T and UC-T controls (21.67 ± 1.16 days for UC-T; 23.33 ± 1.53 days for VC-T; 58.00 ± 4.58 days for T; p<0.05) (Figure 3A). Following six weeks post drug cessation, we found that T males produced consistently smaller litters across three consecutive breedings (Figure 3B). However, although initial litter sizes were reduced from JQ1-treated sires compared to vehicle-treated and untreated sires, pup numbers gradually normalized, reaching comparable levels across all condition groups by the third litter (Figure 3C). Thus, while fertility is eventually restored following JQ1 treatment, the reduced pup number in the early recovery period may reflect compromised sperm quality or quantity. This prolonged recovery of fertility indicates that the health of repopulating spermatozoa may require more time to fully recover following meiosis disruption.

**Figure 3.**
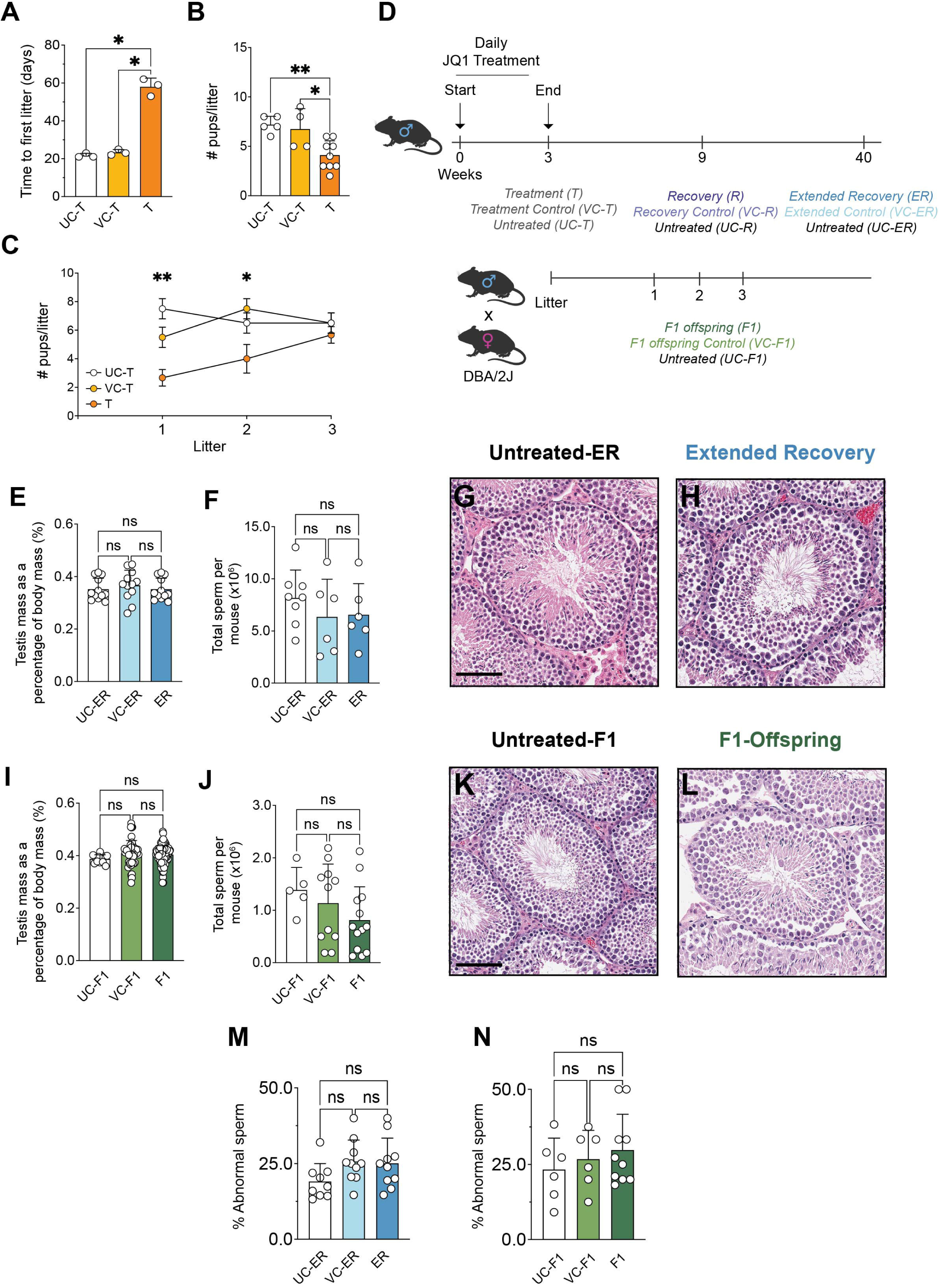
Fertility and sperm abnormalities recover long-term following spermatogenesis resumption. (A) Time to first litter in days from UC-T, VC-T, and T sires (*n* = 3). (B) Number of pups produced per litter from *n* = 3 consecutive litters of UC-T, VC-T, and T sires (*n* = 3 sires). (C) Number of pups per litter across first, second and third litters from UC-T, VC-T and T sires (*n* = 3). Statistics compared UC-T to T, and UC-T to VC-T for each litter number. (D) Experimental design outlining breeding assay and condition groups for extended recovery and F1 generation. After JQ1 or vehicle treatment, males were placed in a rotating breeding assay until siring three litters. Sires from JQ1 (ER), vehicle (VC-ER), and untreated controls (UC-ER) were sacrificed 30 weeks post-treatment for extended recovery analysis. Male offspring from JQ1 (F1), vehicle (VC-F1), and untreated controls (UC-F1) were sacrificed at 7 weeks for F1 analysis. (E) Testis mass per mouse as percentage of body mass (% ± SD) for UC-ER (*n* = 6), VC-ER (*n* = 6), ER (*n* = 6). (F) Total epididymal sperm counts (sperm per million in ± SD) for UC-ER (*n* = 8), VC-ER (*n* = 6), ER (*n* = 6). (G, H) Testicular histology for Untreated-ER (G) and Extended Recovery (H) as shown by H&E staining of paraffin embedded testis sections. (I) Testis mass per mouse as percentage of body mass (% ± SD) for UC-F1 (*n* = 6), VC-F1 (*n* = 25), F1 (*n* = 54). (J) Total epididymal sperm counts (sperm per million in ± SD) for UC-F1 (*n* = 5), VC-F1 (*n* = 11), F1 (*n* = 13). (K, L) Testicular histology for Untreated-F1 (K) and F1 Offspring (L) as shown by H&E. (M, N) Quantification of sperm abnormalities (mean ± SD) from UC-ER (*n* = 9), VC-ER (*n* = 10) and ER (*n* = 10) males, and from UC-F1 (*n* = 6), VC-F1 (*n* = 6) and F1 (*n* = 10) males. Scale bar = 100 µm. P values obtained from a one-way ANOVA test (ns = not significant, *p < 0.05, **p<0.01).

Given the delayed return to normal litter sizes following JQ1 withdrawal, we expanded our experimental timeline to profile spermatogenesis at several stages after withdrawal of JQ1: 6 weeks post drug cessation (Recovery; R), and 30 weeks post drug cessation (Extended Recovery; ER), as well as in the F1 offspring from the first three litters (F1 offspring; F1). This latter group of males were never themselves exposed to JQ1 but were sired from male mice treated with JQ1 (Figure 3D), allowing us to test the transgenerational impact of JQ1 treatment. Our analysis revealed that testis mass, sperm counts, and testicular morphology were normal at both ER and F1 stages (Figure 3E-L). Importantly, sperm from both ER and F1 animals did not exhibit increased abnormalities compared to controls, indicating that sperm quality improves alongside the restoration of fertility for treated mice, and is not impacted in the male offspring of treated mice (19.10 ± 5.89 for UC-ER; 25.67% ± 7.10 for VC-ER; 25.09% ± 8.32 for ER) (Figure 3M) (23.29% ± 10.50 for UC-F1; 26.78% ± 9.58 for VC-F1; 29.82% ± 11.90 for F1) (Figure 3N). Collectively, these data reveal that while sperm production resumes relatively quickly after meiotic disruption, full recovery of sperm quality and functionality takes more time, highlighting the importance of monitoring long-term effects on sperm quality and fertility across generations to evaluate the impact of such contraceptives.

### JQ1 disrupts meiotic progression but recovery precedes full fertility restoration

Studies presented here (Figure 2) show that JQ1 treatment disrupts the transcriptional profile of spermatocytes, an effect that persists for at least six weeks after drug cessation. While germ cells repopulate within this period, the sustained disruption in the spermatocyte transcriptome raises a key question: is meiotic prophase I integrity compromised? Prophase I is critical for spermatogenesis, as it is the main window for transcriptional activation in meiosis, and precise regulation here is essential for producing functional haploid gametes. Importantly, for contraceptives to be truly reversible, fertility restoration must ensure the recovery of all processes affecting gamete production and quality. Thus, we examined whether meiotic prophase I progression is affected upon JQ1 treatment.

To assess meiotic progression, we staged spermatocytes using prophase chromosome spreads stained with SYCP3 to visualize the synaptonemal complex, and γH2AX, which localizes to DNA double strand breaks (DSBs) during early prophase and the sex body domain during pachynema and diplonema (Fernandez-Capetillo et al. 2003; Gray and Cohen 2016) (Figure 4A). In JQ1-treated mice, we observed a significant reduction in pachytene- and diplotene-stage cells compared to controls, indicating that JQ1 disrupts progression through prophase I (Figure 4C). The timing of prophase I disruption is consistent with the onset of BRDT presence, which was observed in the pachynema to diplonema stages of prophase I (Figure 4B). Following withdrawal of JQ1 and recovery in the R group of males, meiotic cells had fully repopulated (Figure 4C), consistent with single-cell RNA-seq data (Figure 2.1H). Thus, the loss of prophase I cells caused by JQ1 treatment, and their subsequent recovery following drug withdrawal, can be observed both by scRNA-seq and by chromosome spreading.

**Figure 4.**
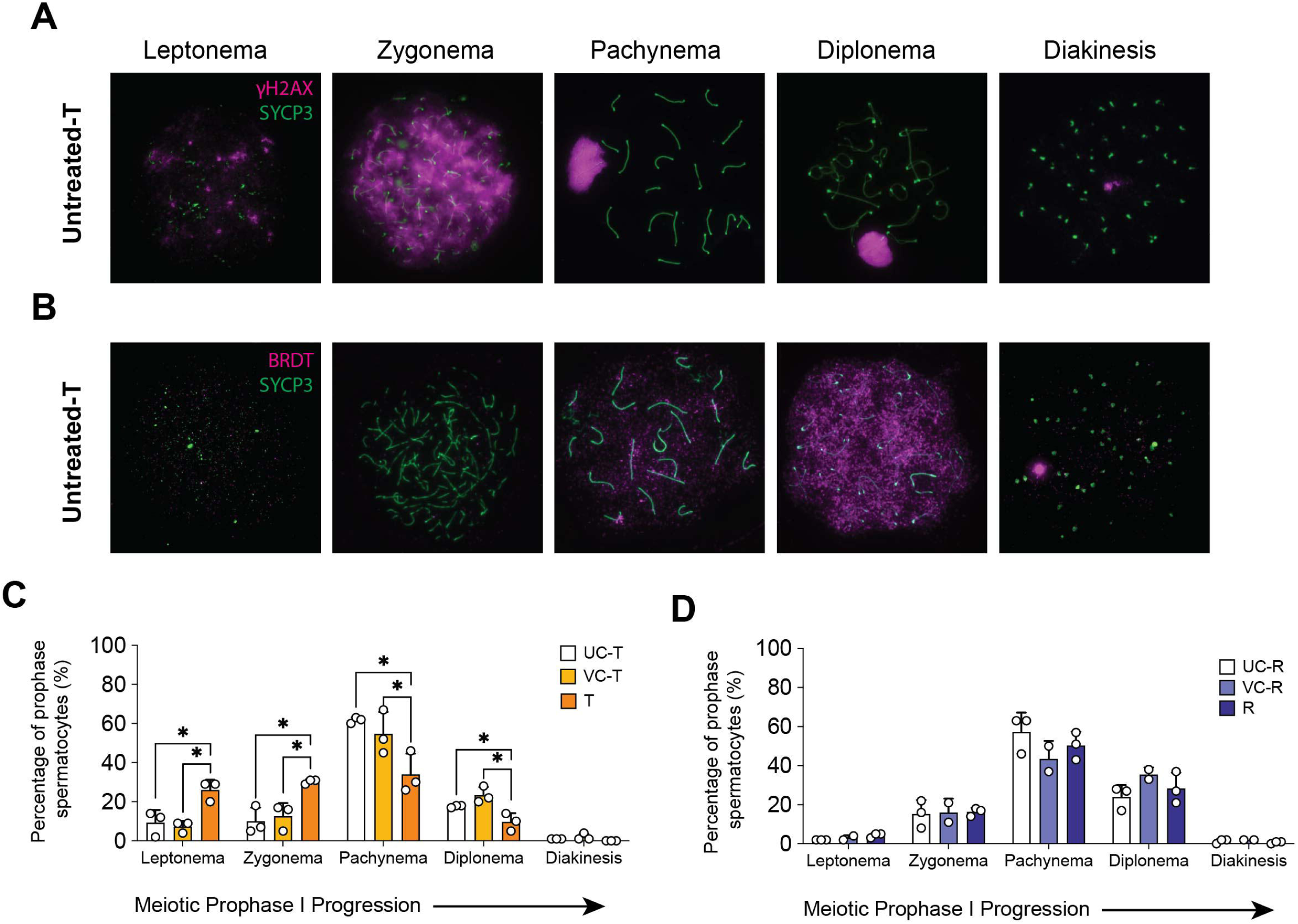
Meiotic prophase I progression is disrupted at the onset of BRDT presence. (A) Prophase I spermatocytes from Untreated-T immunostained with SYCP3 (green) and γH2AX (magenta) showing representative images of the five substages of prophase I – leptonema, zygonema, pachynema, diplonema and diakinesis. (B) Prophase I spermatocytes from Untreated- T immunostained with SYCP3 (green) and BRDT (magenta). (C, D) Percentage of cells (± SD) in each substage of prophase I in (C) UC-T, VC-T and T mice, and (D) UC-R, VC-R and R mice (*n* = 2 or 3 mice per condition group, 100 cells assessed per mouse). P values obtained from a one-way ANOVA test (ns = not significant, *p < 0.05).

### Crossover numbers remain reduced in repopulating spermatocytes, but normalize in ER

Defects in meiotic progression induced by JQ1 treatment, particularly the loss of pachytene and diplotene-staged cells, indicate dysregulation of prophase I, a critical stage where even minor disruptions can have severe downstream consequences for spermatogenesis. Crossover (CO) regulation during prophase I must be precisely controlled to ensure proper chromosome segregation and promote genetic diversity, both of which are essential for gamete quality and fertility (Gray and Cohen 2016). In line with this critical function of COs, any disruption in the frequency or distribution of COs lead to aneuploidy and cell loss. Thus, we were asked whether JQ1 treatment could cause defects in recombination leading to CO formation.

To assess crossover formation, we stained prophase spreads for MLH1, a marker of late-stage recombination (Edelmann et al., 1996). Spermatocytes from JQ1-treated males exhibited significantly fewer MLH1 foci compared to controls (20.41 ± 1.59 for UC-T; 20.38 ± 1.61 for VC-T; 18.35 ± 2.87 for T; p<0.05) (Figure 5A-C). Even modest reductions in CO formation increase the risk of achiasmate chromosomes – homologs that fail to undergo meiotic recombination and consequently segregate improperly (Singh et al. 2021). Consistent with this, spermatocytes from treated males had significantly higher percentage of cells with MLH1 focus-deficient chromosomes (28.57 ± 8.69 for UC-T; 35.42 ± 0.18 for VC-T; 62.88 ± 3.28 for T; p<0.05) (Figure 5D) and fewer chiasmata at metaphase I (23.18 ± 1.33 for UC-T; 22.77 ± 1.74 for VC-T; 21.07 ± 1.03 for T; p<0.05) (Figure 5E-G). However, the percentage of cells with complete bivalents remained largely unchanged (Figure 5H) (100 ± 0.00 for UC-T; 98.89 ± 1.92 for VC-T; 93.86 ± 5.90 for T; p<0.05), suggesting that while CO formation is impaired, homolog synapsis is still achieved. These findings indicate that JQ1 treatment reduces CO events, increasing the risk of achiasmate chromosomes and altered recombination patterns.

**Figure 5.**
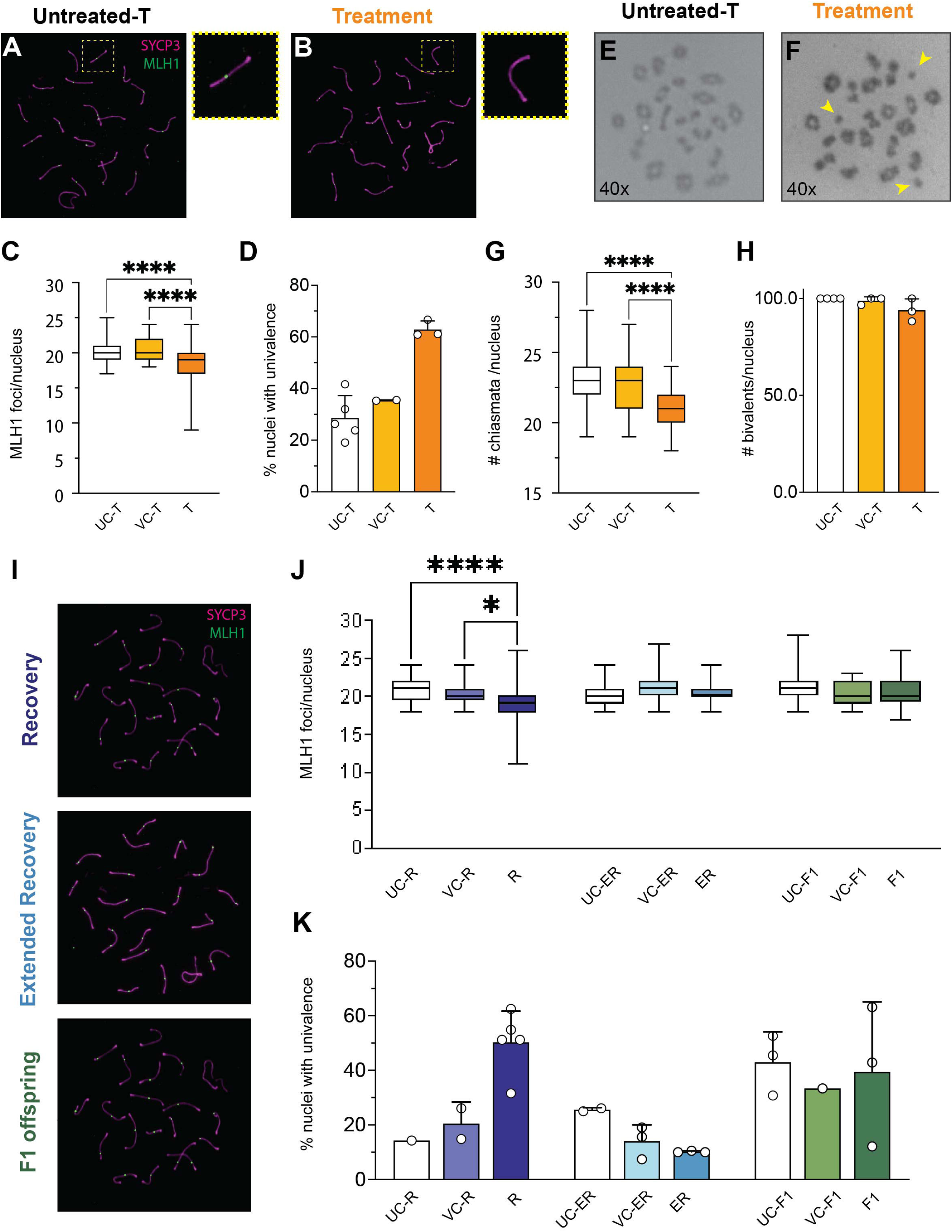
Disrupted CO numbers normalize by extended recovery. (A, B) Pachytene spermatocytes from Untreated-T (A) and Treatment (B) mice immunostained with antibodies to MLH1 (green) and SYCP3 (magenta). Yellow dashed box indicate zoomed image of a single SC either with (A) or without (B) MLH1 focus. (C) Comparison of mean MLH1 foci number for UC-T (*n* = 5), VC-T (*n* = 3), and T (*n* = 4) males. (D) Comparison of mean percentage of cells with univalence from UC-T (*n* = 5), VC-T (*n* = 2), and T (*n* = 3) males. (E, F) Chiasmata stained with Giemsa from Untreated-T (E) and Treatment (F) males. Yellow arrows indicate univalence. (G) Comparison of mean chiasmata counts for UC-T (*n* = 5), VC-T (*n* = 3) and T (*n* = 4) males. (H) Percentage of cells with bivalents per nuclei from UC-T (*n* = 4), VC-T (*n* = 3) and T (*n* = 3) males. (I) Pachytene spermatocytes from recovery (R), extended recovery, (ER) and F1 mice immunostained with antibodies to MLH1 (green) and SYCP3 (magenta). (J) Comparison of mean MLH1 foci number for UC-R (*n* = 3), VC-R (*n* = 3), R (*n* = 5), UC-ER (*n* = 4), VC-ER (*n* = 3), ER (*n* = 3), UC-F1 (*n* = 4), VC-F1 (*n* = 3), and F1 (*n* = 4) males. (K) Comparison of mean percentage of cells with univalence from UC-R (*n* = 1), VC-R (*n* = 2), R (*n* = 5), UC-ER (*n* = 2), VC-ER (*n* = 3), ER (*n* = 3), UC-F1 (*n* = 3), VC-F1 (*n* = 1), and F1 (*n* = 3) males. At least 10 cells analyzed per mouse for each condition group. Data represent the mean ± SD and are annotated with P values obtained from a one-way ANOVA test (ns = not significant, *p < 0.05, ****p<0.0001).

Following recovery and withdrawal of JQ1, CO formation remained impaired, with MLH1 foci counts in R spermatocytes still significantly reduced compared to VC-R and UC-R controls and similar to those observed in T spermatocytes (20.75 ± 1.47 for UC-T; 20.30 ± 1.22 for VC-T; 19.24 ± 2.27 for T; p<0.05) (Figure 5I-J). The frequency of MLH1 focus-deficient chromosomes remained elevated (14.29 ± 0.00 for UC-R; 20.45 ± 7.97 for VC-R; 50.25 ± 11.42 for R; p<0.05) (Figure 5K), and metaphase I chiasmata counts did not recover (Supplementary Figure S7), indicating that defects in late-stage recombination persisted after six weeks of drug withdrawal. However, following 30 weeks post drug cessation (ER), we observed recovery in CO numbers, with MLH1 foci counts (20.32 ± 1.38 for UC-ER; 21.00 ± 1.93 for VC-ER; 20.43 ± 1.23 for ER; p<0.05) (Figure 5J) and chiasmata numbers (Supplementary Figure S7) approaching those of controls. CO dynamics were excitingly unperturbed in the F1 spermatocytes, where MLH1 focus counts (20.81 ± 1.79 for UC-F1; 20.45 ± 1.41 for VC-F1; 20.84 ± 1.92 for F1; p<0.05) and chiasmata (Supplementary Figure S7) were comparable across all three groups (Figure 5J, K).

Taken together with our breeding data and sperm analysis, these results indicate that withdrawal of the JQ1 treatment does not fully recover molecular parameters of prophase I and sperm functionality within 6 weeks post-drug cessation. Since meiotic prophase I is one of the longest phases of spermatogenesis, taking up to 13 days in mouse and at least double that time in humans (Griswold 2016), the full recovery of prophase I fidelity is likely to be time-consuming, which may account for the persistence of defects during this period. However, the complete recovery of these processes following extended recovery, with no defects carried through into the next generation, provides strong evidence supporting the potential of meiotic prophase I as a target for contraceptive intervention.

## DISCUSSION

The development of male-targeted contraceptives presents an opportunity to alleviate the burden of family planning unduly placed on women, while equalizing the balance of decision-making between men and women in efforts to reduce the high rates of unintended pregnancies (Bearak et al. 2020). However, a major factor limiting the development of non-hormonal contraceptive options for males is the relative absence of data to assist in identifying suitable intervention points for contraception. The current study was designed to test whether meiotic prophase I represents such a suitable intervention point by disrupting mid-prophase I with the BRDT inhibitor JQ1. Moreover, this study systematically examined the molecular dynamics of spermatogenic recovery across a timecourse designed to reflect the duration and reversibility required for translational relevance to human male contraception. Daily injections of JQ1 for three weeks led to disrupted meiotic prophase I progression (Figure 4), CO dynamics such as fewer MLH1 numbers and chiasmata (Figure 5), and altered transcription of spermatocytes (Figure 2), ultimately resulting in a loss of post-meiotic spermatids and spermatozoa and conferment of a contraceptive effect (Figure 1). Following six weeks post drug withdrawal, we observed a repopulation of post-meiotic cells within the seminiferous tubules; however, the transcriptional burst in spermatocytes remain altered even after six weeks, consistent with the presence of abnormal spermatozoa and aberrant CO numbers. This observation prompted us to extend our recovery assessment to thirty weeks post drug cessation and to examine male offspring sired by JQ1-treated animals. Full recovery of recombination events was observed after thirty weeks of drug withdrawal, with similar results seen in males sired by JQ1-treated animals (Figures 4, 5). Our findings demonstrate that the small molecule inhibitor JQ1 disrupts meiosis by altering the pachytene transcriptional burst, leading to a loss in post-meiotic cells. Importantly, these effects are conclusively reversible, supporting the pursuit of meiotic prophase I as a suitable intervention target for non-hormonal male contraception. These results have crucial implications for considering targets during meiotic prophase I for male contraception, opening up avenues for future contraceptive strategies that act by disrupting key events during meiosis.

Efforts to develop non-hormonal male contraception have primarily targeted either the earliest or latest stages of spermatogenesis. YCT529, for example, disrupts chromatin relaxation and gene expression in undifferentiated spermatogonia by inhibiting retinoic acid receptor alpha (Abdullah Al Noman et al. 2020). In contrast, mature sperm-targeting approaches include EP055, which blocks the prostate-specific antigen by binding semenogelin (SEMG1) (O’Rand and Widgren 2012; O’Rand et al. 2018), and soluble adenylyl cyclase (sAC), which facilitates capacitation by generating cyclic AMP in response to calcium and bicarbonate in semen (Balbach et al. 2023). While promising, these strategies would likely necessitate daily intake or provide only short-term, on-demand contraception. Nevertheless, no studies have explored the potential of targeting meiotic processes for contraception, nor have they assessed the reversibility of meiotic disruption – highlighting a critical gap in the development of effective and reversible non-hormonal male contraceptives.

To date, no studies – including those studying the effects of JQ1 or other BRDT inhibitors – have investigated the effects of reversible disruption of meiotic prophase I as a potential intervention stage for contraception (Filippakopoulos et al. 2010; Matzuk et al. 2012; Wang et al. 2022; Alexander et al. 2023). Our study addresses this gap by demonstrating that meiotic prophase I can serve as a viable target for non-hormonal male contraception, with a clear loss of spermatozoa following three weeks of treatment. One intriguing observation from our study is the discrepancy following drug cessation. While spermatozoa are repopulated within six weeks, abnormalities in the molecular regulation during pachynema persist, including features of the transcriptional burst that is characteristic of this prophase I substage (Figure 2). Furthermore, the repopulating spermatozoa exhibit persistent morphological abnormalities (Figure 2) which impact fertility, since litter numbers only normalize by the third litter sired by JQ1-treated males. These findings underscore the importance of thoroughly evaluating spermatogenesis when considering meiosis as a target for male contraception. Since meiosis represents a pivotal checkpoint in germ cell development, disruptions to meiosis can lead to widespread alterations in transcriptional regulation, sperm maturation, and overall fertility. This necessitates a comprehensive investigation across all stages of spermatogenesis.

Despite the availability of several BRDT inhibitors (Yu et al. 2021; Modukuri et al. 2022), we chose the small-molecule inhibitor JQ1 as our contraceptive drug to disrupt meiotic prophase I (Matzuk et al. 2012). JQ1 is commercially available, and its contraceptive effects have been well characterized, allowing us to follow the *in vivo* dosing regimen and timeline previously outline (Matzuk et al. 2012). This gave us the opportunity to extend the recovery timelines and analyses beyond fertility endpoints, enabling us to draw specific conclusions on the effects of prophase I disruptions within the broader context of spermatogenesis. Much is already known about the mechanism and specificity of JQ1. It effectively targets BRDT and is classified as a pan-BET inhibitor, with varying affinities for binding to the three other members of the BET family of proteins (Matzuk et al. 2012). *Brdt* is testis-specific in both mouse (Shang et al. 2004) and human (Jones et al. 1997), suggesting that any effects from BRDT inhibition are likely confined to the testis. However, other BET family members, including *Brd4*, are also expressed during spermatogenesis, with BRD4 being important for spermatogenesis progression and expressed earlier than BRDT (Berkovits and Wolgemuth 2013). Therefore, while BRDT inhibition is a primary target, we cannot rule out the possibility that BRD4 inhibition may also contribute to the observed phenotype.

While JQ1 treatment elicits a contraceptive effect, its phenotypic consequences do not fully align with those observed in *Brdt* knockout models. To date, both a full *Brdt*-null allele (*Brdt*^-/-^) as well as a *Brdt*-null allele lacking the first bromodomain (*Brdt*^ΔBD1/ΔBD1^) have been generated and characterized. Although both mouse mutants are sterile, *Brdt*^-/-^ male mice exhibit a complete loss of post-meiotic cells and a mid-prophase I arrest (Shang et al. 2004; Manterola et al. 2018). In contrast, *Brdt*^ΔBD1/ΔBD1^ mice complete meiosis and produce spermatozoa, but these sperm display aberrant sperm head morphology, leading to complete sterility (Shang et al. 2007). Since JQ1 competitively binds to the bromodomains of BRDT (Matzuk et al. 2012), we anticipated that its phenotypic effects would resemble those of the *Brdt*^-/-^ mutant mice. While we observed disruptions in meiotic prophase I progression and chromosomal features (Figures 4, 5), the extent of disruption was far less severe than in *Brdt*^-/-^ mutant mice (Manterola et al. 2018). This discrepancy may be due to the transient inhibition of BRDT by JQ1, as opposed to the permanent loss of function seen in genetic knockouts. Furthermore, the expression of other BET proteins during spermatogenesis may compensate for the inhibition of BRDT, lessening the severity of the JQ1 effects (Shang et al. 2004). As research on contraceptive strategies increasingly targets meiotic prophase I, understanding the differences between pharmacological inhibition and genetic knockouts will become crucial. Our findings suggest that targeting essential meiotic regulators, such as BRDT, results in persistent disruptions in CO formation and spermiogenesis. We know from previous studies that complete loss of CO proteins like MLH1 leads to meiotic arrest (Jackson et al. 1996). Furthermore, recombination defects that disrupt bivalent formation in only a few chromosome pairs are often insufficient to activate the spindle assembly checkpoint, allowing these gametes to persist and increasing the risk of aneuploidy (Singh et al. 2021). Therefore, it will be important to investigate whether small-molecule inhibitors of specific meiotic regulators produce similar discrepancies and lead to an increased risk of aneuploid gametes upon drug cessation. Addressing these questions will be essential for refining drug-based approaches to male contraception.

Given the human translational focus of this study, an important question arises: How does a three-week treatment duration in a murine model translate to a human contraceptive timeline? Previous studies in the United States have shown that over 80% of women use oral contraceptives, such as the pill, for an average of five years during their reproductive years (Tyrer 1999). Based on these estimates, it seems reasonable to suggest that a male contraceptive, once available, could be used for a similarly long duration, or potentially longer, due to the extended reproductive span of males. While short-term treatment regimens in mammalian models targeting one spermatogenic cycle offer valuable insights, it remains uncertain whether these results can fully extend to human contraceptive timelines. Specifically, we observed a discrepancy between the timeline of cell repopulation and the reversibility of meiotic disruptions, with spermatozoa repopulating within six weeks but continuing to show abnormalities. This suggests that longer treatment durations and more extensive evaluations of reversibility at key stages will be necessary. Future studies should consider extending treatment durations to better align with typical human contraceptive use and carefully evaluate the reversibility of effects at these key stages before advancing to human trials.

Taken together, the data presented here demonstrates that meiotic prophase I can be effectively targeted for male contraception, with pharmacological disruption proving to be reversible. While treatment leads to initial disruptions in meiotic progression and cell loss during spermiogenesis, these effects are not permanent, as repopulating cells gradually recover over time. However, some abnormalities in meiosis such as CO dynamics and aberrations in sperm maturation persist, highlighting the complexity of reversibility. These findings are significant in establishing meiosis as a viable target for male contraception and provide a critical foundation for future research aimed at testing drug targets within this stage of spermatogenesis. This study offers an important step toward developing targeted contraceptive strategies that focus on meiotic prophase I.

## ACKNOWLEDGEMENTS

We are grateful to Cornell Center for Animal Resources and Education (CARE) for assistance with mouse husbandry and Eileen Shu for overall lab management as well as the Cohen lab for feedback and advice during the course of this research. We thank Peter Schweitzer and colleagues in the Cornell Biotechnology Resource Center for their help with preparing the Chromium datasets.

## Author contributions

Conceptualization: L.E.S.*, S.T.*, P.E.C., C.G.D., A.K.A., J.L.

Methodology: L.E.S.*, S.T.*, P.E.C., A.K.A., J.L., C.G.D.

Software: S.T., C.G.D., A.K.A.

Formal analysis: L.E.S.*, S.T.*, P.E.C., C.G.D., S.J.B., R.B.S., A.X.

Investigation: L.E.S.*, S.T.*, J.L., T.S.H., M.C., S.J.B.

Resources and data curation: L.E.S.*, S.T.*, C.G.D.

Funding acquisition: P.E.C.

Writing (original draft): L.E.S.*, S.T.*, P.E.C.

Writing—review and editing: L.E.S.*, S.T.*, P.E.C., C.G.D., J.L.

*: Authors contributed equally

## Data and materials availability

All data are available in the main text.

## FIGURE LEGENDS

**Supplemental Figure S1. Preprocessing and quality control metrics of scRNA-seq data.** (A) Workflow used for preparation, integration, and analysis of scRNA-seq compendium, compartmentalized by analysis performed on individual samples and merged datasets. (B) Violin plots showing the quality control for the scRNA-seq data, displaying the distribution of gene counts, UMI counts, and mitochondrial gene percentage per library before filtering data. The red dash lines represent quality control process standard. (C) Principal component analysis (PCA) of spermatogenic cells based on scRNA-seq dataset. Condition groups are presented in distinct colors, and each point corresponds to a distinct library per condition group. (D) Pearson correlation coefficient between filtered cells separated by condition groups using log-normalized average expression of all genes.

**Supplemental Figure S2. Genes used as markers of somatic cells captured in scRNA-seq dataset.** UMAP plots showing representative markers for somatic cells: Sertoli cells (*Cst9, Rhox8*); Leydig cells (*Cyp17a1, Hsd3b1*); Myoid cells (*Acta2, Timp3*); Macrophages (*Pf4, Lyz2*).

**Supplemental Figure S3. Genes used as markers to define germ cells captured in the scRNA-seq dataset.** UMAP plots showing representative markers for spermatogonia (*Sall4, Dmrt1*), spermatocyte (*Sycp3, Shcbp1l*), spermatid (*Acrv1, Ccna1*), and spermatozoa (*Oaz3, Prm2*).

**Supplemental Figure S4. Increased trends of apoptotic cells observed following 3 weeks JQ1 treatment.** TUNEL assay indicating apoptotic cells as shown by DAB staining, co-stained with Methyl Green, followed by quantification of DAB positive cells across per tubules across entire testis section for (A) UC-T (*n* = 5), VC-T (*n* = 6) and T (*n* = 6) males, and (B) UC-R (*n* = 5), VC-R (*n* = 3) and R (*n* = 4) males. Scale bar = 100 µm. P values obtained from one-way ANOVA test (ns = not significant).

**Supplemental Figure S5. Pseudotime trajectory analysis of germ cells from scRNA-seq dataset.** (A) Pseudotime trajectory analysis along UMAP of germ cells from scRNA-seq. Arrow indicates the differentiation order of germ cells from spermatogonia to spermatozoa. (B) Line graphs showing relative expression patterns of representative marker genes of each germ cell type along pseudotime. Note different Y axis scales.

**Supplemental Figure S6. BRDT levels in spermatocytes are unaffected following JQ1 treatment.** (A) UMAP visualization of all cells captured following Untreated control from scRNA-seq showing *Brdt* expression. (B, C) Violin plots showing *Brdt* mRNA expression in spermatocytes (B) and spermatids (C) in individual cells, grouped by condition. Statistical comparisons are based on average expression per biological replicate (*n* = 3 per group). P values obtained from a one-way ANOVA test (ns = not significant). (D-F) Immunofluorescence staining of paraffin-embedded testis sections with DAPI (greyscale) and BRDT (magenta) showing presence of BRDT in (D) UC-T, (E) VC-T and (F) T males.

**Supplemental Figure S7. Chiasmata numbers recover following 30 weeks post drug cessation.** Comparison of mean chiasmata counts 6 weeks post drug cessation: UC-R (*n* = 4), VC-R (*n* = 3) and R (*n* = 5) males; 30 weeks post drug cessation: UC-ER (*n* = 3), VC-ER (*n* = 3) and ER (*n* = 3) males; F1 generation: UC-F1 (*n* = 4), VC-F1 (*n* = 2) and F1 (*n* = 4) males. Data represent the mean ± SD and are annotated with P values obtained from a one-way ANOVA test (ns = not significant, *p < 0.05, **p<0.01, ***p<0.001).

## MATERIALS AND METHODS

### Study design

The goal of this study was to assess the suitability of meiotic prophase I as an intervention stage for non-hormonal male contraceptive testing. To do this, we used the small-molecule BRDT inhibitor JQ1 as a tool to induce blockade of meiotic prophase I by injecting mice daily for three weeks with the drug. Following this, we assessed for spermatogenesis disruption using scRNA-seq, histology, and chromosome spreading. As this was a contraceptive study, we assessed recovery following drug cessation for spermatogenesis progression at six weeks, and thirty weeks post drug cessation. Finally, we assessed spermatogenesis in the F1 males sired from JQ1-treated males. A minimum of three biological replicates (individual mice) per group were used, based on precedent in all of our previous studies. All data were unbiasedly collected and analyzed in single-blind studies. The findings in this study were collected from multiple independent experiments and were reliably reproduced. The *n* numbers of each sample are indicated in the figure legend.

### Mice

The experiments described used mice on the DBA/2J backgrounds, obtained from Jackson Laboratories (Strain #000671). All male mice were 7 weeks old at the start of experiments. Female mice used for breeding were 8 weeks old when introduced with a male. Male mice at 7, 10, 15, and 34 weeks of age, used as aged-matched, untreated controls, were also obtained from Jackson Laboratories. All mice were housed under strictly controlled conditions of temperature and light:day cycles, with food and water *ad libitum*. All mouse studies were conducted with prior approval by the Cornell Institutional Animal Care and Use Committee, under protocol 2004-0063.

### JQ1 dosage and injections

JQ1 dosage followed the original study by (Matzuk et al. 2012). Briefly, JQ1 sourced from AdooQ Bioscience (CAS # 1268524-70-4) was dissolved in DMSO at an initial concentration of 50 mg/ml and then diluted 1:10 in 10% (2-Hydroxypropyl)-β-cyclodextrin (Sigma Aldrich, H5784). This mixture was injected intraperitoneally (i.p.) into male mice at 1% body weight for 3 weeks. Vehicle control animals received a similar daily i.p. injection of DMSO dissolved 1:10 in 10% (2-Hydroxypropyl)-β-cyclodextrin. All injected mice were weighed daily prior to injections, and doses were adjusted accordingly.

### Breeding assay

Male mice (*n* = 3) were used for breeding either immediately following 3 weeks JQ1 treatment or vehicle treatment. Ten-week-old male mice from Jackson Laboratories were used as the untreated control. Eight-week-old DBA/2J females were placed in a paired breeding assay for 14 days with each of the males. New females were placed in breeding every 14 days until each male sired a total of three litters. Throughout the breeding duration, sires did not receive any treatment. Male offspring from each litter were weaned upon sexual maturity and housed under the same conditions previously stated until analysis at 7 weeks of age.

### Single cell testicular suspension

One testis was collected for single cell RNA sequencing (*n* = 3 mice). Testes were dissociated as described previously (Grive et al. 2019) with minor modifications. Briefly, testes were detunicated and minced on ice and transferred to a 50 mL tube containing 40 ml of collagenase IV (1 mg/ml, Sigma-Aldrich, C4-BIOC) and DNase I (7 mg/mL, Sigma Aldrich DN25) in Dulbecco’s Modified Eagle Medium (DMEM, Gibco, 11-965-092). Tissue was incubated in a 37 °C incubator for 15 minutes. Tubule fragments were then centrifuged at 60 x g for 10 minutes, the supernatant discarded, washed with 15 mL Dulbecco’s Phosphate Buffered Saline (dPBS, Sigma-Aldrich, D8537), and centrifuged again at 60 x g for 10 minutes. The fragments were then incubated in 10 mL 0.05% trypsin-EDTA (Invitrogen, 25200056) solution in 37 °C incubator for 12 minutes. Trypsin was quenched with 10% v/v fetal bovine serum (FBS, Thermo Fisher Scientific, A5256701), cells were passed through a 40 µm filter, spun down at 300 x g for 5 minutes, and resuspended in 1mL 0.04% bovine serum albumin (BSA, Sigma-Aldrich, A4737) in 1X dPBS. The resulting cells from all samples were submitted to the Cornell DNA Sequencing Core Facility for further processing.

### Single cell RNA sequencing

We performed three-prime (3’) Chromium 10X Genomics cell capture and cDNA library preparation at each timepoint, aiming to capture approximately 8,000 cells per library (*n* = 3 mice per condition group). The resulting 15 libraries were pooled at equal concentrations and sequenced across two lanes of a NovaSeq flow cell (Novogene, California, USA) to minimize batch effects. Details on estimated sequencing saturation and the number of reads per sample are provided in Supplementary Table 1.

### Preprocessing and quality control of single-cell RNA sequencing datasets

Data processing and quality control were performed to ensure high-confidence single-cell transcriptomic data (Figure S1A). The ‘cellranger count’ pipeline (v3.0.0) was used with default parameters to process raw sequencing data, and the STAR aligner (v2.5.3b) mapped reads to the mm10 reference genome (downloaded from the 10X Genomics website).To minimize background noise, ambient RNA signal was first removed from each dataset using the default SoupX (v1.4.5) workflow (github.com/constantAmateur/SoupX). Each sample was then individually pre-processed using the standard Seurat (v4.1.1) workflow, including NormalizeData, ScaleData, FindVariableFeatures, RunPCA, FindNeighbors, FindClusters, and RunUMAP (github.com/satijalab/Seurat). Low-quality cells were filtered out by removing those with fewer than 500 features, fewer than 1,000 transcripts, or more than 30% of unique transcripts derived from mitochondrial genes (Figure S1B). To further refine cell quality, DoubletFinder (v2.0) was applied to identify and remove putative doublets, with estimated doublet rates computed based on 10X Chromium guidelines. Following quality control, individual datasets were merged for further processing using Seurat.

A single treatment control sample (TC1) showed unexpected similarity to Treatment samples in PCA space and was flagged as a potential outlier (Figure S1C). This observation was reinforced by pairwise Pearson correlation analysis (Figure S1D), where Treatment Control showed reduced correlation with the other control conditions (r = 0.93–0.94), and its similarity to Treatment samples inflated the apparent correlation between Treatment and Treatment Control groups (r = 0.51). Based on this evidence, TC1 was retained for unsupervised analyses (e.g., UMAP, pseudotime) but excluded from condition-level comparisons during differential expression testing to avoid skewing results.

### Cell type annotation

Cell types were assigned based on the expression of well-established canonical marker genes (Figure S2, 3). Clusters exhibiting highly similar marker gene expression patterns were merged to refine classification. A clustering resolution of 1.2 was applied, initially generating 32 clusters. Following the merging of biologically similar clusters, we identified five major cell types and 21 sub-clusters, representing all stages of spermatogenesis (Figure S3).

### Pseudotime workflow

Pseudotime trajectory analysis was performed to infer the developmental progression of germ cells through spermatogenesis. The merged Seurat object was first subset to include only germ cells and then analyzed using the default Slingshot (v2.4.0) workflow (Street et al. 2018). UMAP embeddings were extracted and used as the reduced dimensionality input. Lineage trajectories were inferred using cluster labels, and pseudotime values were extracted using the slingPseudotime function. Lineage trajectories were overlaid using the SlingshotDataSet function. Cells were binned into 51 pseudotime intervals to facilitate the analysis of gene expression dynamics across spermatogenesis.

### Differential expression analysis

To identify differentially expressed genes between conditions, we performed bulk-like differential expression analysis using Seurat’s AggregateExpression function. Gene expression was aggregated at the sample level, grouping by condition and sample identity. Differential expression analysis was conducted using the Seurat FindMarkers function with the DESeq2 framework (Love et al. 2014). Differentially expressed genes were filtered based on a false discovery rate (FDR)-corrected *P* value threshold of *p* < 0.05. Additionally, genes were selected if they had an average log_2_-fold change (avg_logFC > 0.5) and were expressed in all samples (pct.expressed = 1). Due to the presence of an outlier sample in the Treatment Control (TC) group, we limited our differential comparisons to UC vs. Treatment and UC vs. Recovery to ensure accurate representation of condition-specific effects.

### Sperm capture methods

#### Total epididymal sperm collection

The caudal epididymides of untreated mice and mice treated with JQ1, or vehicle control were transferred and dissected into sperm count Toyoda, Yokoma, Hoshi (TYH) media (Toyoda and Yokoyama 2016). The samples were incubated at 37°C for 20 minutes in order to allow sperm to swim out into the media. Dilutions of each sample were made to a 1:10 concentration in 10% neutral buffered formalin. Spermatozoa were counted and analyzed for morphological defects using a hemocytometer.

#### Hematoxylin-eosin staining of epididymal sperm smears

Ten microliters of total epididymal sperm were added to 10 μl of 4% paraformaldehyde (PFA), spread on a glass slide, and air dried for 6 hours at room temperature. Slides were stored at 4°C until use. Slides were washed in PBS and subject to Hematoxylin and Eosin staining using standard methods.

### Prophase I chromosome spreading and immunofluorescence staining

Prophase I chromosome spreads and immunofluorescent staining were prepared as previously described by (Holloway et al. 2014) with minor modifications. Testis tubules were incubated in hypertonic elution buffer (30 mM Tris-HCl pH 7.2, 50 mM sucrose, 17 mM trisodium dehydrate, 5 mM EDTA, 0.5 mM DTT, 0.1 mM PMSF, pH 8.2-8.4) for 20 minutes. Small sections of tubules were minced in 100 mM sucrose solution and spread onto 1% Paraformaldehyde, 0.15% Triton X-100 coated slides. Slides were incubated in a humid chamber for 2 hours at room temperature. Slides were dried for at least 30 minutes, washed in 0.4% Photoflo (KODAK, 1464510) diluted in PBS (800 µL in 200 mL PBS), 0.1% Triton X-100 (Fisher Scientific, BP151100) diluted in PBS (1 mL and 199 mL PBS) and blocked in 10% antibody dilution buffer (ADB: 3% BSA, 1% Goat serum [Gibco, 16210-072], 0.05% Triton in 1X PBS) diluted in PBS (20 mL and 180 mL). Primary antibodies used included rabbit-anti-SYCP3 1:1,000 (custom-made; (Kolas et al. 2005)), mouse anti-γH2AX 1:10,000 (EMD Millipore, 05-636), mouse-anti-SYCP3 1:1,000 (Abcam, ab97627), mouse-anti-MLH1 1:50 (BD Pharmingen, 550838), and rabbit-anti-BRDT 1:500 (Abcam, ab286194). Primary antibodies were diluted in ADB, spread across the surface of the slide, and incubated at room temperature overnight. Slides were washed for 10 minutes each in 0.4% Photoflo, 0.1% Triton X, and 10% antibody dilution buffer. Alexafluor™ secondary antibodies (Molecular Probes Eugen) were used for immunofluorescent staining at room temperature for 2 hours. Secondary antibodies were diluted in antibody dilution buffer and spread in a similar fashion to the primary antibodies. Slides were washed as previously described, dried, and mounted with Prolong Gold antifade (Molecular Probes) with DAPI. All secondary antibodies were from Jackson ImmunoResearch and were diluted 1:1000, including: AffiniPure F(ab’)_2_ fragment goat anti-mouse IgG Fc fragment specific conjugated to AF488 (115-546-008) or AF594 (115-586-008); anti-rabbit IgG Fc fragment specific conjugated to AF488 (111-546-046) or AF594 (111-586-046). Slides were stored at 4 °C prior to analysis.

### Diakinesis preparations to observe chiasmata

Diakinesis chromosome preparations were prepared as described previously with slight modifications (Holloway et al. 2010). Briefly, testes were detunicated and seminiferous tubules were macerated in 2.2% sodium citrate. Tubules were further dissociated by pipetting and larger fragments of tubules were left to settle for 15 minutes. The supernatant cell suspension was collected and centrifuged for 5 minutes at 600 x g. The resulting cell pellet was resuspended in 4 mL 1.1% sodium citrate and incubated at room temperature for 15 minutes. The cell suspension was centrifuged for 5 min at 600 x g and the sodium citrate supernatant was discarded. The cell pellet was vigorously vortexed into suspension as fixative (3:1 methanol:glacial acetic acid) was slowly added to the cells. Following 2 additional centrifugations and resuspension in fixative, cells were dropped on slides and air dried. Slides were stained for 6 minutes with 4% Giemsa (Fisher), washed three times for 3 minutes each with ddH_2_O, dried, and mounted with permount.

### Histological methods

#### Sample collection

One testis was sampled from each mouse in all condition groups and fixed in 10% formalin overnight at room temperature, following which they were washed for 30 minutes each in 30%, 50% and then 70% ethanol. Paraffin-embedded tissue was sectioned at 5 µm and processed for Hematoxylin and Eosin staining or immunohistochemical analyses using standard methods.

#### TUNEL staining of testicular sections

TUNEL staining was performed using a TUNEL staining kit (Apoptag kit, EMD Millipore, S7100) following the manufacturer’s instructions. Images were obtained using Image Scope, and DAB positive cells were manually counted by Adobe Photoshop software. Number of apoptotic cells was assessed by counting the total number of DAB positive cells, then dividing by the total number of tubules per each testis section.

#### Immunohistochemistry of testicular sections

Immunohistochemistry was performed as described previously (Holloway et al. 2010; Holloway et al. 2011; Holloway et al. 2014). Briefly, slides were rehydrated in Safe Clear Xylene Substitute and ethanol. Sections were permeabilized using 0.1% Triton-X for 15 minutes. Antigen retrieval was performed by boiling the slides in sodium citrate buffer (10 mM sodium citrate, 0.05% Tween 20, pH 6.0) for 25 minutes. Following subsequent cooling, slides were blocked in blocking buffer (1X PBST, 1% BSA, 3% goat serum) for an hour and primary antibody dilutions were incubated on the sections for an hour at 37°C. Antibodies used included mouse-anti-SYCP3 1:1000 (as stated above), rabbit-anti-BRDT 1:500 and PNA-lectin 1:500 (GeneTex). Slides were washed and incubated with fluorescence-conjugated secondary antibodies for one hour at 37°C, washed and then mounted using a DAPI/Antifade mix. Slides were imaged on a Zeiss Axiophot with Zen 2.0 software. All secondary antibodies were used at 1:500. For assessment of staging, seminiferous tubules were split into categories (I, II-III, IV–VI, VII–VIII, IX-X, XI-XII) according to their lectin staining pattern and cell types present, as previously described (Nakata et al. 2015).

### Image acquisition

Images were acquired using a Zeiss Axiophot Z1 microscope attached to a cooled charge-coupled device (CCD) Black and White Camera (Zeiss McM). The images were captured and pseudo-colored by means of ZEN 2 software (Carl Zeiss AG, Oberkochen. Germany). Brightness and contrast of images were adjusted using ImageJ (National Institutes of Health, USA) and Adobe Photoshop. Exposure time was consistent between antibodies, cells, and test groups.

### Statistical analysis

Statistical analyses were completed using GraphPad Prism version 6.0 for Macintosh (GraphPad Software, San Diego California USA, www.graphpad.com). Kruskal-Wallace tests or One-way ANOVAs were performed to determine statistical significance. Mean values were presented ± standard deviation (SD) and alpha values were established at 0.05. All statistical analyses performed utilized two-sided tests.

**Table S1.** Metadata for datasets included in this study.

| Sample Name | Condition | Number of reads | Reads mapped | Sequencing saturation | Estimated cell # prior to filtering | Estimated cell # after filtering | Cell # after Doublet Removal |
| --- | --- | --- | --- | --- | --- | --- | --- |
| JQ3WR1 | Recovery | 243,444,103 | 94.6% | 32.0% | 9,949 | 6,408 | 6,055 |
| JQ3WR2 | Recovery | 275,053,459 | 90.4% | 32.6% | 10,785 | 7,191 | 6,792 |
| JQ3WR3 | Recovery | 439,265,185 | 91.0% | 34.7% | 14,221 | 12,739 | 12,031 |
| JQ3WRC1 | Recovery Control | 152,539,066 | 94.6% | 28.1% | 6,740 | 4,898 | 4,634 |
| JQ3WRC2 | Recovery Control | 170,172,586 | 96.0% | 35.2% | 5,218 | 3,592 | 3,401 |
| JQ3WRC3 | Recovery Control | 97,538,675 | 96.4% | 31.3% | 3,407 | 2,836 | 2,684 |
| JQ3WT1 | Treatment | 175,844,767 | 95.2% | 28.8% | 9,567 | 5,694 | 5,551 |
| JQ3WT2 | Treatment | 161,894,445 | 95.2% | 34.0% | 4,787 | 3,974 | 3,763 |
| JQ3WT3 | Treatment | 144,378,699 | 94.9% | 32.5% | 3,573 | 2,527 | 2,397 |
| JQ3WTC1 | Treatment Control | 155,707,767 | 96.0% | 30.5% | 6,932 | 5,746 | 5,443 |
| JQ3WTC2 | Treatment Control | 200,624,393 | 96.2% | 36.5% | 7,080 | 3,521 | 3,330 |
| JQ3WTC3 | Treatment Control | 167,287,369 | 96.0% | 34.5% | 5,042 | 3,595 | 3,405 |
| UC1 | Untreated | 149,774,138 | 96.1% | 34.3% | 4,149 | 3,160 | 2,993 |
| UC2 | Untreated | 154,707,378 | 92.9% | 38.3% | 4,071 | 2,975 | 2,817 |
| UC3 | Untreated | 167,822,360 | 96.2% | 35.0% | 6,089 | 4,191 | 3,966 |
|  |  | **Total:** |  |  | **69,292** |  |  |

