## Supplementary figures and images for "Meiotic prophase I disruption as a strategy for non-hormonal male contraception using small-molecule inhibitor JQ1"

### Supplemental Figure 1

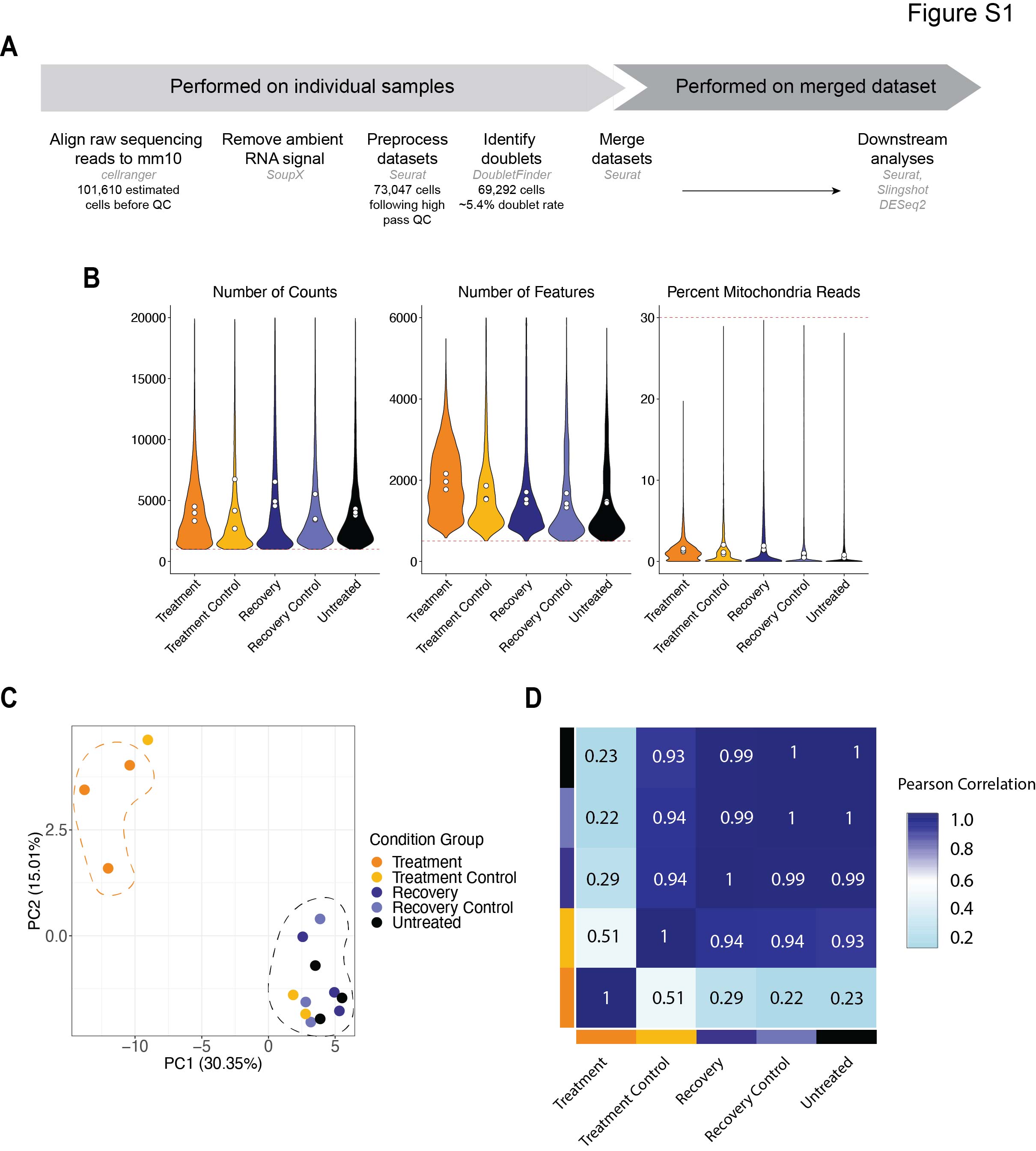

### Supplemental Figure 2

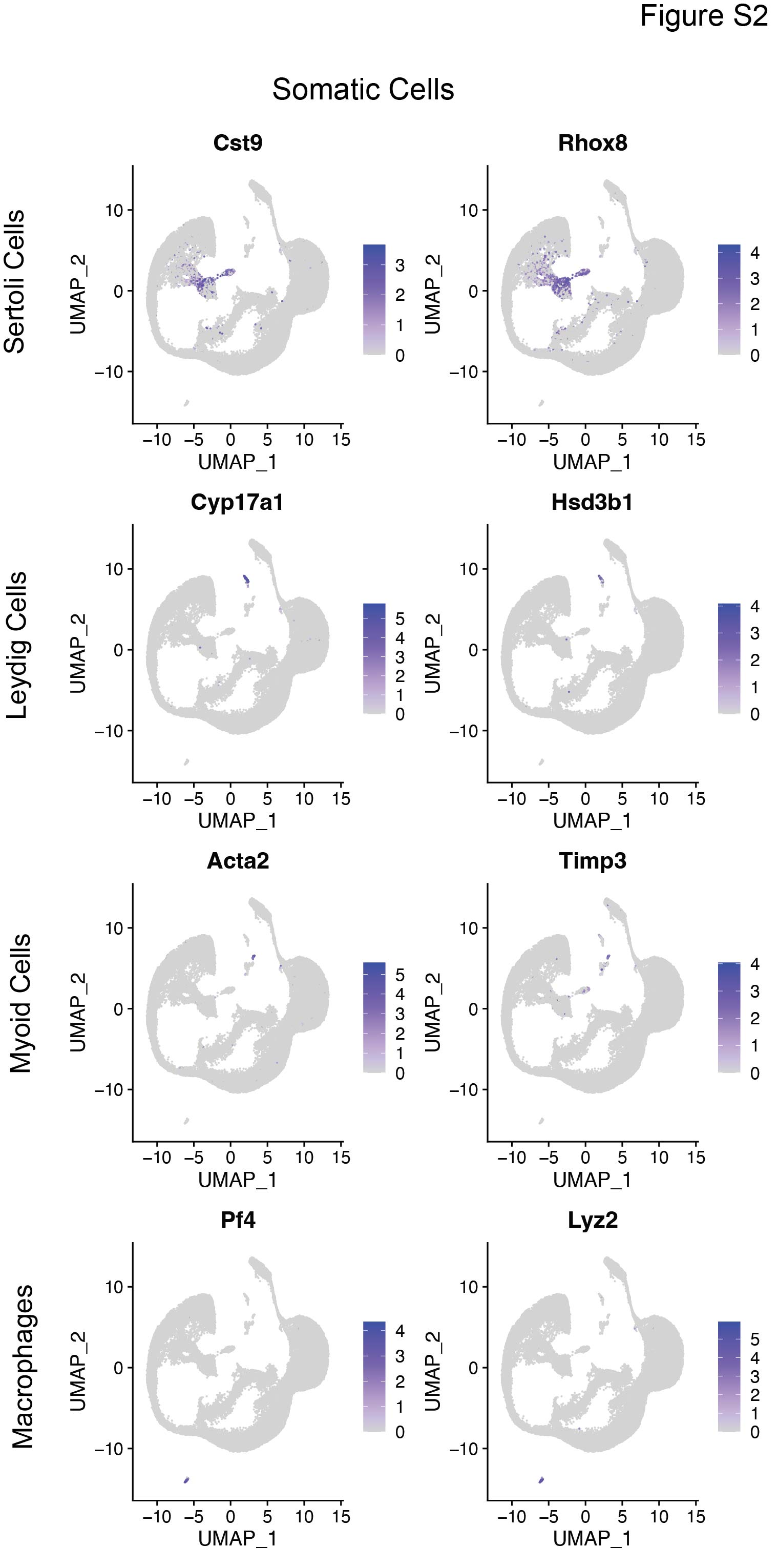

### Supplemental Figure 3

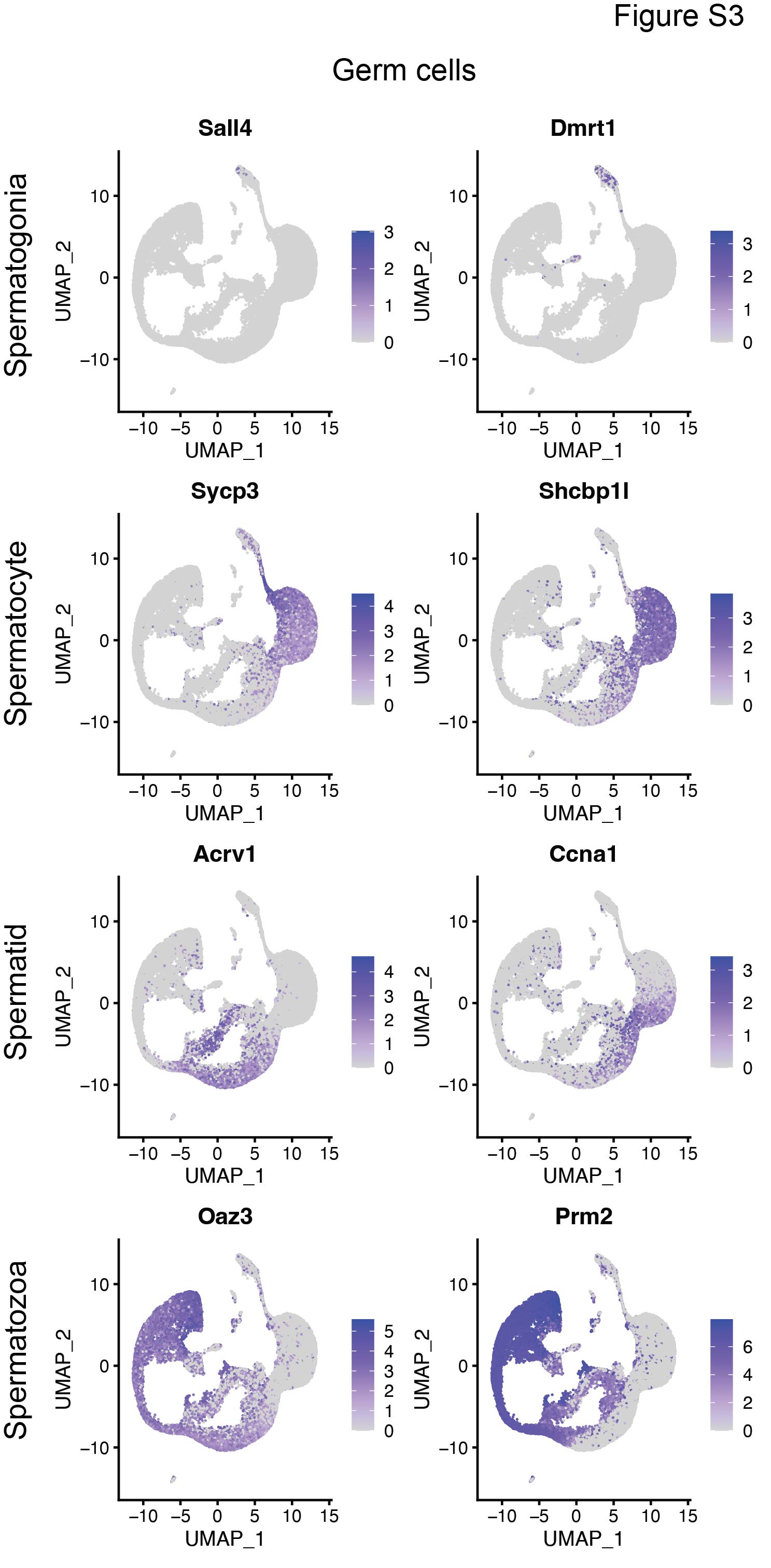

### Supplemental Figure 4

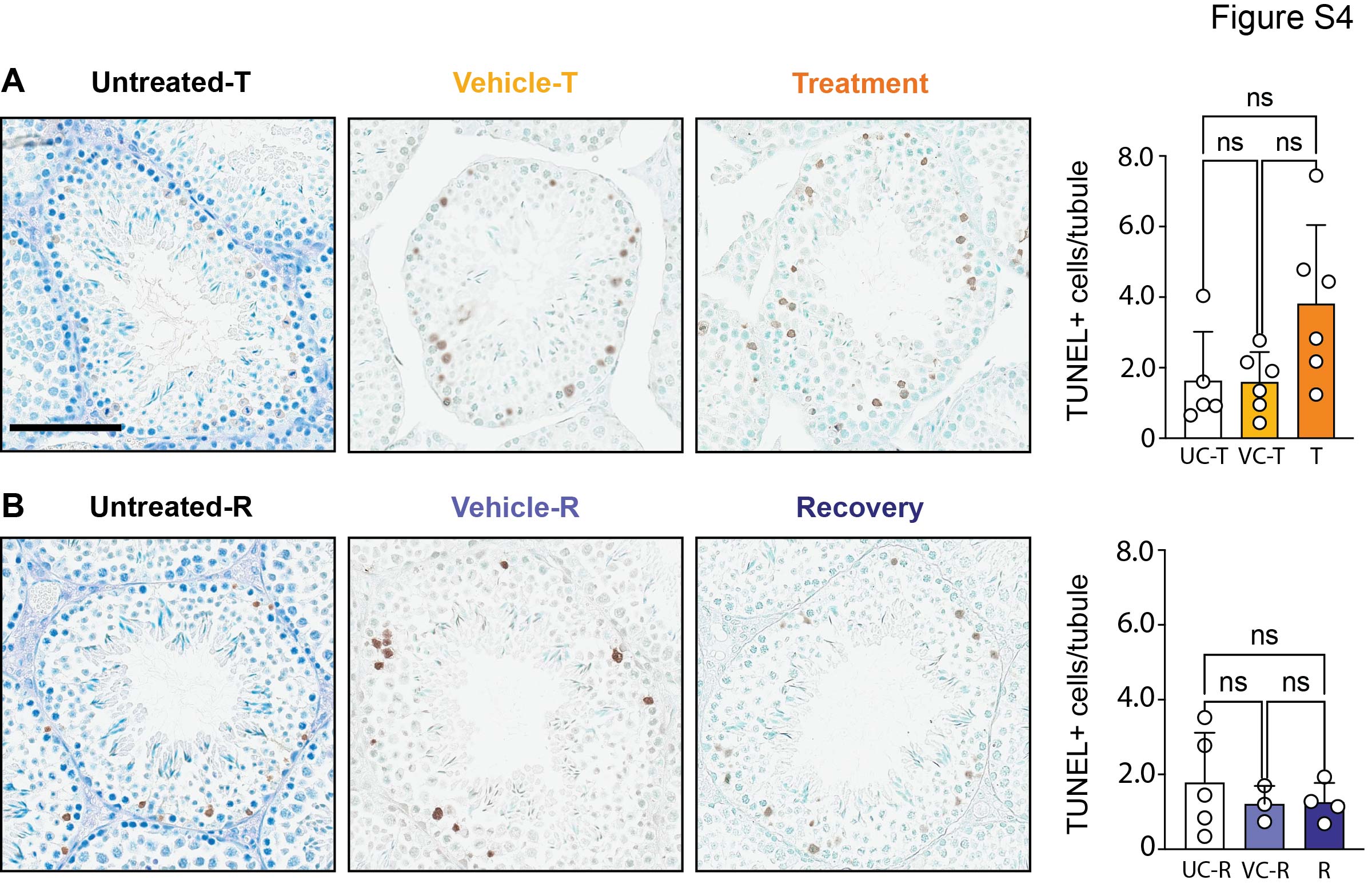

### Supplemental Figure 5

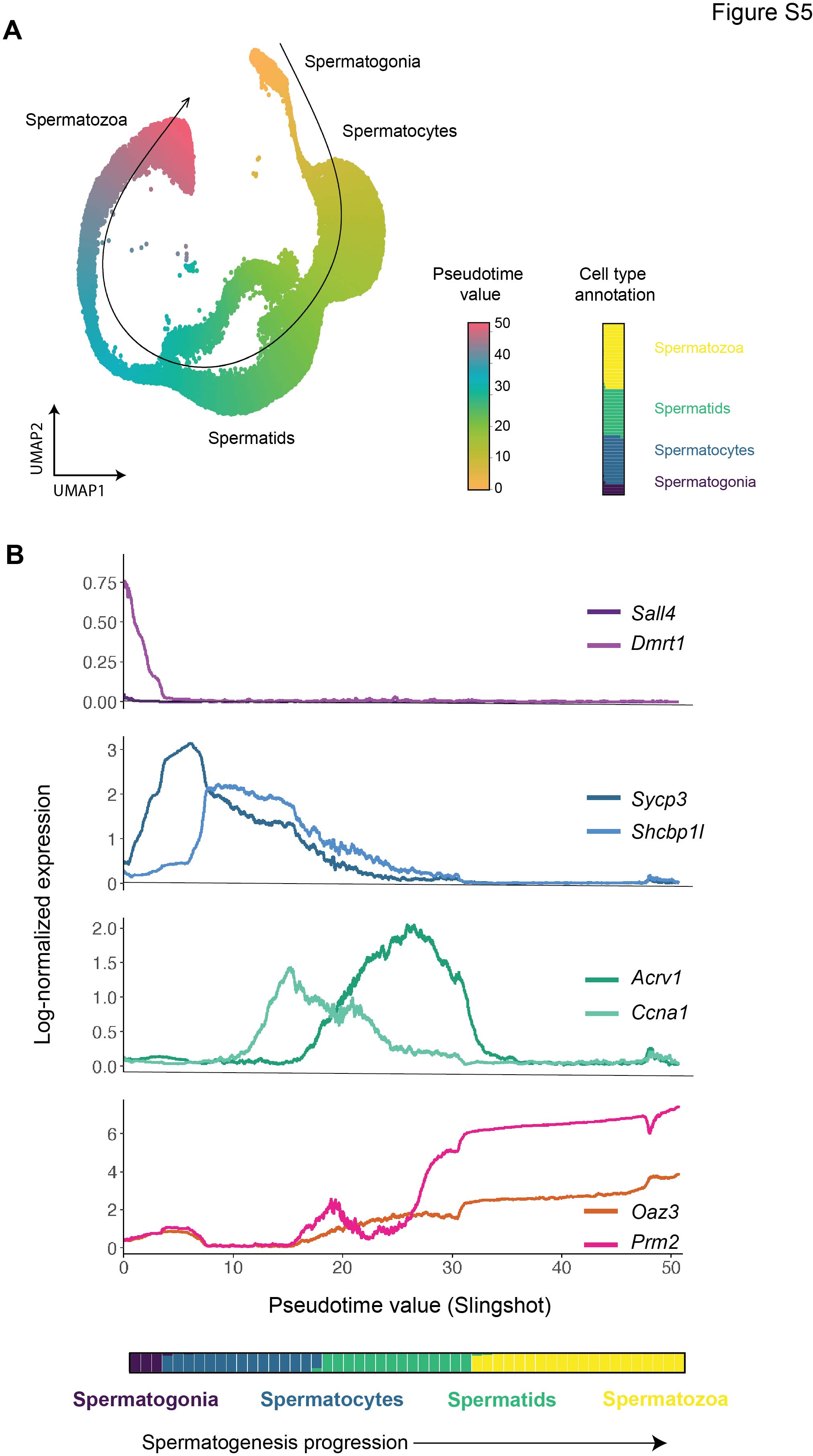

### Supplemental Figure 6

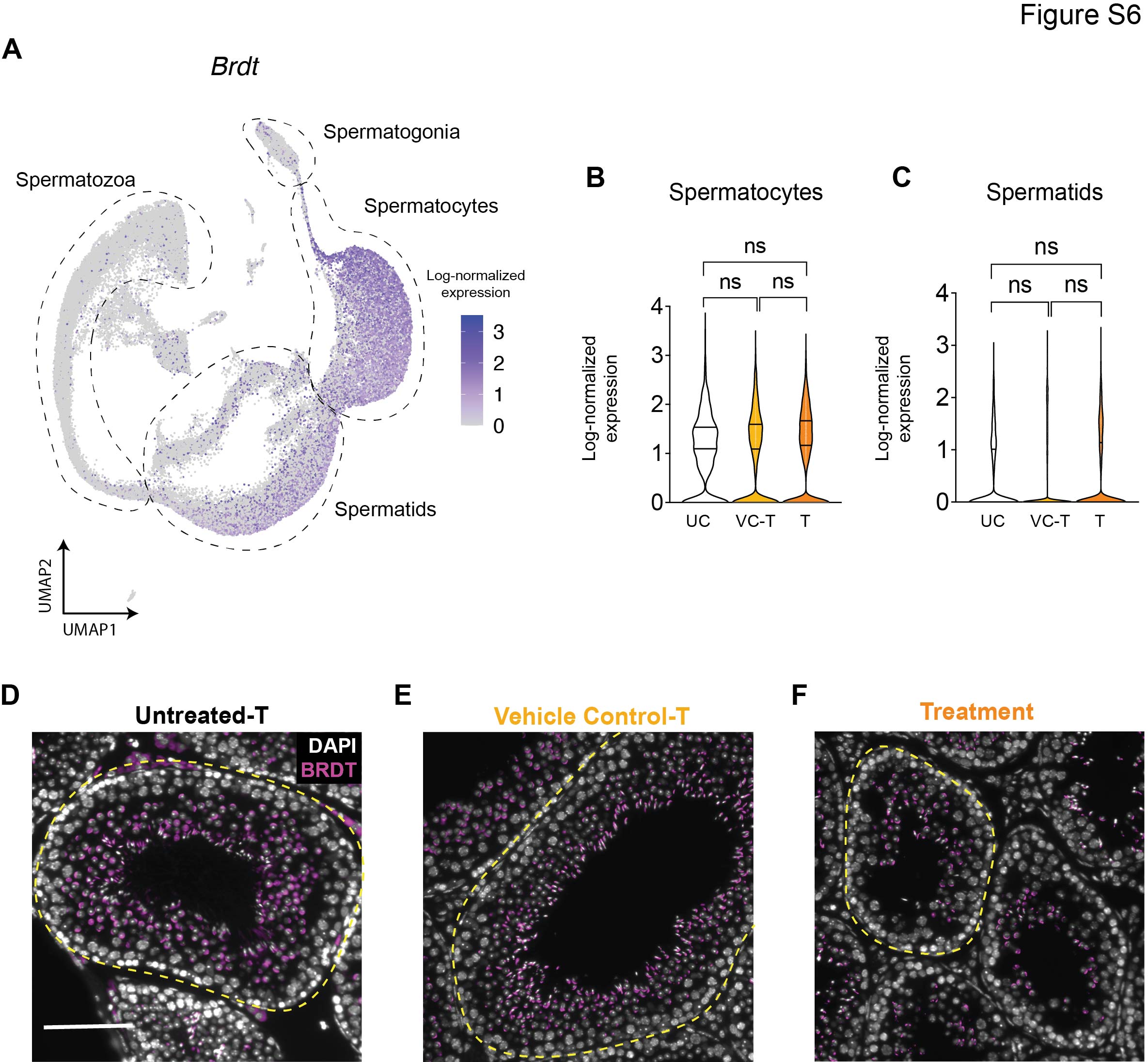

### Supplemental Figure 7

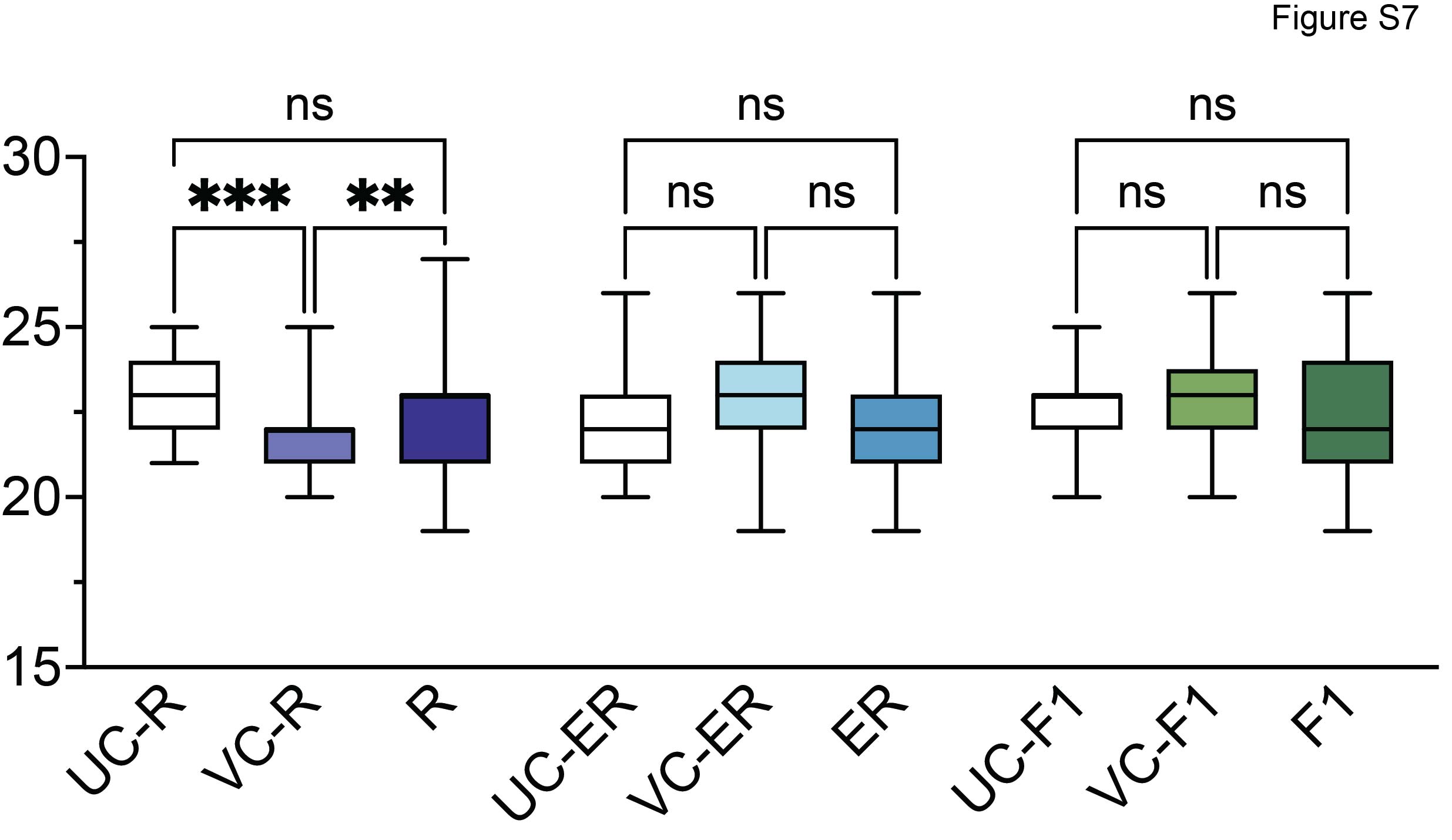
